# HLF suppresses DREAM target genes and reinstates quiescence in lung squamous cell carcinoma

**DOI:** 10.1101/2025.05.06.650933

**Authors:** Pavan Prabhala, Sofia Freiman, Jiten Sharma, Julia Stevrell, Sofia Wijk, Frida Rosengren, Camila Vazquez Echegaray, Nika Gvazava, Karina Kanzenbach, Stefan Lang, Rebecka Cattani, Jenny Wigen, Shamit Soneji, David Bryder, Elsa Arbajian, Johan Flygare, Ellen Tufvesson, Leif Bjermer, Darcy Wagner, Hans Brunnström, Maria Planck, Gunilla Westergren Thorsson, Mattias Magnusson

## Abstract

Lung squamous cell carcinoma (LUSC) originates from airway basal stem cells and carries frequent TP53 mutations and RB pathway disruption, yet lacks oncogene-directed therapies. Here, using single-cell transcriptomics of healthy airway epithelium and primary LUSC tumors, we resolve four basal cell states along a quiescence-to-activation gradient and identify HLF as a transcription factor restricted to the most primitive quiescent population. In LUSC, HLF is silenced across all six genetically distinct tumors and associated with promoter hypermethylation, and low HLF expression predicts poor patient survival. Concurrent with HLF silencing, DREAM target genes are broadly de-repressed through convergent DREAM and RB1 pathway disruption. Functionally, HLF re-expression reinstates G0 quiescence and suppresses proliferation in basal stem cells, LUSC cell lines, and patient-derived tumor organoids, preferentially suppressing DREAM targets over E2F-only targets. Because HLF engages DREAM target repression independently of p53, reactivation could restore quiescence in LUSC regardless of p53 status.

**Statement of Significance:** We identify convergent DREAM complex disruption as the proliferative mechanism in LUSC and show that the quiescence factor HLF suppresses DREAM target genes independently of p53, reinstating G0 arrest in cancer cells. HLF is silenced by promoter methylation, representing a pharmacologically accessible target amenable to reactivation.

## Introduction

Lung squamous cell carcinoma (LUSC) accounts for approximately 30% of non-small-cell lung cancers and carries a poor prognosis, with five-year survival rates below 20% for late-stage disease (1,2). Unlike lung adenocarcinoma, where targetable driver mutations in EGFR, ALK, and KRAS have transformed treatment, LUSC lacks targetable drivers and is treated primarily with platinum-based chemotherapy and immune checkpoint inhibitors (2,3). Genomic profiling has revealed that the majority of LUSC tumors carry TP53 mutations (∼80%) and harbor alterations in the RB pathway (4). Despite the identification of these frequent alterations, no effective targeted therapies have emerged, underscoring the need for therapeutic approaches at the transcriptional level.

The cell of origin of LUSC is thought to be the airway basal stem cell (BSC), a TP63+/KRT5+ progenitor responsible for lifelong renewal and repair of the airway epithelium (5–8). Lineage tracing and injury-repair studies have demonstrated that KRT5+ basal cells serve as multipotent stem cells capable of regenerating the full airway epithelium (5–8). Single-cell atlases have defined the cellular composition of the human airway in detail highlighted by the Human Lung Cell Atlas (9–12). Additionally, lineage-resolved profiling of the tracheal epithelium has connected epithelial states into differentiation trajectories (13). In line with that, recent functional studies argue that bronchial basal cells form a continuum of transitional states rather than discrete subtypes (14). These resources annotate basal populations by marker expression and differentiation potential, but they do not assign transcription factor (TF) programs to positions along the basal continuum, identify which program sustains the quiescent state, or ask whether these programs are silenced during transformation. In healthy tissue, BSCs are maintained in a quiescent state (G0), from which they can be transiently activated to proliferate before returning to quiescence or committing to differentiation. How this tightly regulated quiescence program is dismantled during malignant transformation remains an open question with direct therapeutic implications.

The quiescence program is governed by two parallel tumor suppressor pathways that control distinct phases of the cell cycle (15–19). The DREAM complex (Dimerization partner, RB-like p130, E2F4, and MuvB) is the master transcriptional repressor of G0. When assembled in its repressive configuration (p130-E2F4-MuvB), it binds CDE/CHR promoter elements and silences over 210 G2/M cell cycle genes (20). DREAM assembly depends on the p53-p21 axis, where p53 induces p21, which inhibits CDK2 and thereby maintains p130 in its hypophosphorylated, DREAM-competent state. When DREAM disassembles, the MuvB core switches partners, recruiting B-MYB and FOXM1 to form the MMB-FOXM1 complex, which activates the same promoters and drives mitotic gene expression (15–18). In parallel, the RB1 pathway controls the G1/S transition. Here, p16 (CDKN2A) inhibits CDK4/6 to prevent RB1 phosphorylation, allowing hypophosphorylated RB1 to restrain E2F1/2/3 and keep G1/S genes silent (18).

In LUSC, both pathways are frequently disrupted. TP53 is mutated in approximately 80% of cases (4), abolishing p21-mediated CDK inhibition and causing DREAM complex disassembly and G2/M gene de-repression. Concurrently, the RB pathway is altered in the majority of tumors, freeing E2F1/2/3 to drive G1/S gene expression (4). CDK4/6 inhibitors, which act through RB1, have failed in LUSC, with palbociclib achieving only a 6% response rate in the Lung-MAP S1400C trial (21). Restoring DREAM-mediated repression has been proposed as a therapeutic strategy but has not yet been demonstrated (22). This highlights the need to determine which transcription factors maintain quiescence in airway basal stem cells, and which of these are lost during squamous transformation.

To systematically identify the transcription factors that maintain quiescence in the airway basal compartment, we map the transcriptionally distinct basal states and test whether factors lost during squamous transformation can restore repression of the DREAM target program. We resolve transcriptionally distinct states within the basal compartment and use pseudotime-based TF mapping to build a transcriptional roadmap of BSC quiescence, differentiation, and proliferation. This approach identifies Hepatic Leukemia Factor (HLF), a PAR bZIP factor that maintains quiescence in hematopoietic stem cells (23,24), as a regulator of quiescence also in airway basal stem cells, where it is silenced across all LUSC tumors. Here, we show that HLF re-expression reinstates G0 quiescence in normal BSCs, LUSC cell lines, and patient-derived tumor organoids, and that HLF selectively suppresses DREAM targets rather than RB1/E2F targets, revealing that a tissue-specific basal cell transcription factor can engage DREAM target repression independently of p53. TCGA analysis identifies promoter hypermethylation as associated with HLF silencing and reveals that HLF loss predicts poor survival.

## Results

### An integrated single-cell atlas of the healthy airway and LUSC

To identify the transcriptional programs distinguishing normal BSCs from their malignant counterparts, we generated a combined single-cell atlas incorporating bronchial biopsies from 10 healthy never-smoker donors (ages 20-75) and tissue resections from 6 LUSC patients (All KRT5+ with no therapeutic intervention before surgery, all >65 years and from different stages (Supplementary Table 1)), yielding 21,860 and 17,447 high-quality cells after CD45 depletion to exclude immune cells (Fig. 1A, Supplementary Fig. E1, E3, E4, Supplementary Table 1, 2). All six tumors were selected independent of genetic background and showed transcriptional evidence of TP53 pathway dysfunction, though the degree and route varied between patients (Supplementary Fig. E7). After quality control, integration, and iterative clustering, UMAP visualization resolved all expected healthy airway epithelial populations (basal, suprabasal, goblet, secretory, multiciliated, and ionocyte), alongside cancer epithelial cells, AT2 cells, myofibroblasts, and endothelial cells from the tumor resection (Fig. 1B, Supplementary Fig. E2, Supplementary Table 3). Cells from cancer patients and healthy donors segregated clearly (Fig. 1C), with cancer epithelial cells expressing basal lineage markers KRT5, TP63 and EPCAM (Fig. 1D, E), consistent with their BSC origin and a block in differentiation. A dot plot of canonical markers confirmed the identity of all populations (Fig. 1E). Gene expression profiling across all populations revealed that cancer epithelial cells were most similar to healthy basal cells (Fig. 1F). Gene ontology analysis confirmed enrichment for p53 signaling, G2M checkpoint, glycolysis, PI3K/AKT/mTOR signaling, and MYC targets in cancer epithelial cells (Fig. 1G, Supplementary Table 4). Healthy BSCs exhibited enrichment for translation-related pathways, including ribosome assembly, cytoplasmic translation, and peptide biosynthesis, compared to all other cell types including cancer (Fig. 1G-I). Individual ribosomal protein large-subunit (RPL) and small-subunit (RPS) gene expression confirmed this pattern across all healthy basal cells but not cancer epithelial cells (Fig. 1I, Supplementary Fig. E5). This translational priming, previously documented in hematopoietic stem cells (25) but unreported in airway BSCs, suggests that BSCs maintain elevated protein-synthetic readiness to enable rapid activation upon demand. Notably, cancer epithelial cells did not retain this ribosomal signature despite their high proliferative rate, suggesting that translational priming is a feature of the quiescent basal state rather than of proliferation itself. Together, our integrated atlas confirms BSC origin for LUSC, reveals coordinated activation of proliferative and metabolic programs across genetically distinct tumors, and identifies translational priming as a distinguishing feature of normal BSCs.

**Figure 1.**
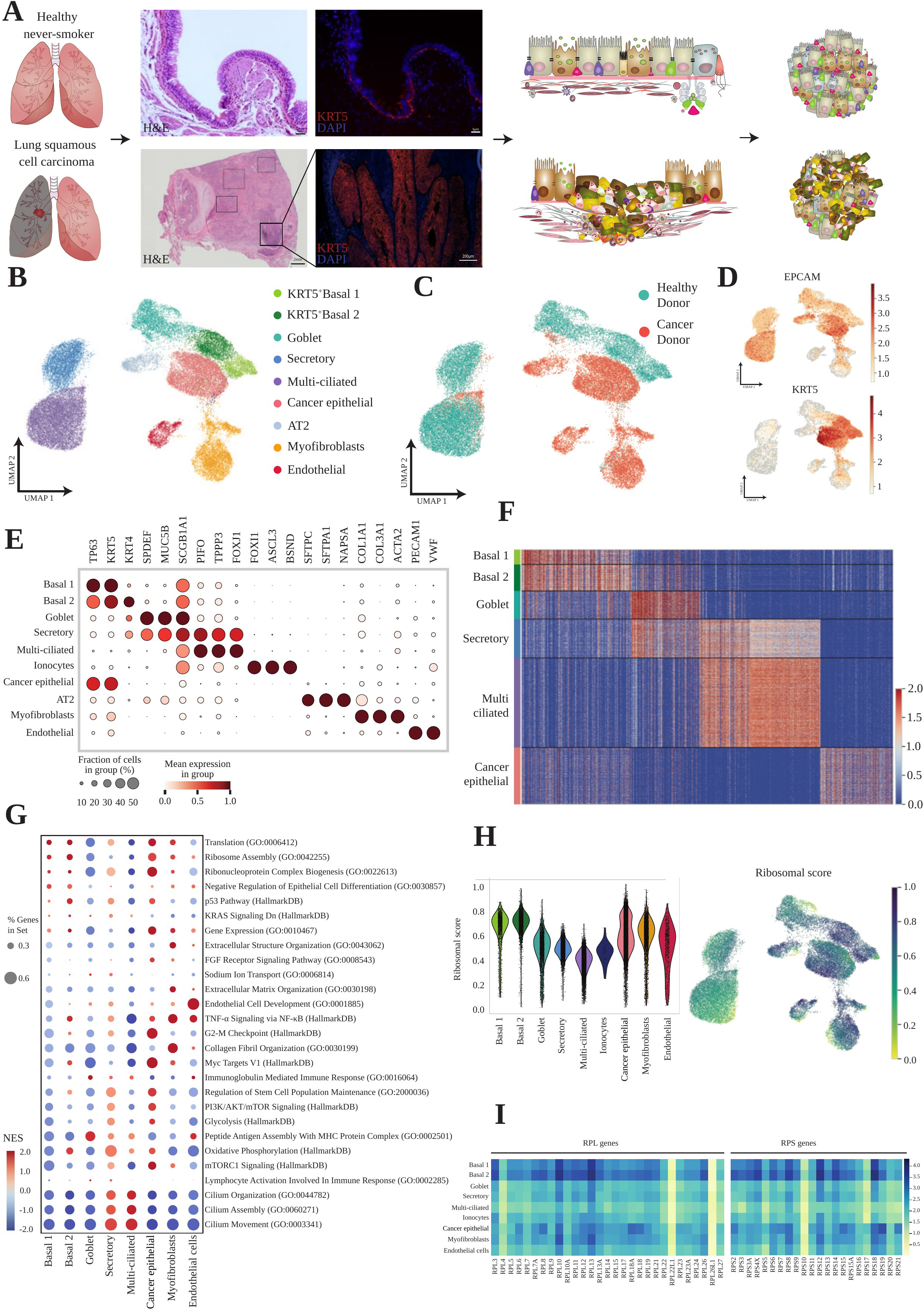
An integrated single cell atlas of healthy airway and LUSC. A, Overview of the study design, showing representative hematoxylin and eosin staining and immunofluorescence for KRT5 and DAPI in a healthy never-smoker airway and in a lung squamous cell carcinoma tumor, together with a schematic of the workflow used to generate single cell RNA sequencing data from normal and tumor tissue. B, UMAP plot of all profiled cells colored by the nine annotated cell types, including KRT5-positive Basal 1 and Basal 2, Goblet, Secretory, Multi-ciliated, Ionocyte, Cancer epithelial, AT2, Myofibroblast, and Endothelial populations. C, The same UMAP embedding colored by tissue origin, distinguishing cells from healthy donors and cancer donors. D, UMAP feature plots showing normalized expression of the epithelial marker EPCAM and the basal marker KRT5. E, Dot plot of canonical marker genes used to annotate each cell type, including Ionocytes, with dot size representing the fraction of cells expressing the gene and color representing mean expression within the group. F, Heatmap of single cell marker gene expression across Basal 1, Basal 2, Goblet, Secretory, Multi-ciliated, and Cancer epithelial cells, with individual cells ordered within each cell type and colored by scaled expression. G, Dot plot of gene set enrichment results across cell types for selected Gene Ontology, Hallmark, and curated pathways, with dot size indicating the percentage of genes in the set detected and color indicating normalized enrichment score. H, Violin plots of a ribosomal gene score across the nine cell types, together with UMAP plots colored by this ribosomal score. I, Curated heatmap of individual large subunit and small subunit ribosomal protein gene expression across all annotated cell types.

### LUSC exhibits convergent disruption of the DREAM and RB1 quiescence axes

Having established that cancer epithelial cells are transcriptionally most similar to healthy basal cells, we next asked what molecular programs distinguish these populations. Cancer cells showed highest correlation with healthy basal cells (R² = 0.60–0.67, Fig. 2A), consistent with a BSC origin. Despite the genetic heterogeneity among the six tumors, we focused on identifying shared transcriptional programs rather than patient-specific alterations. Differential expression analysis identified 399 genes significantly upregulated and 875 downregulated (Supplementary Table 5 and Table 12) in cancer versus basal cells (padj < 0.05, |log2FC| > 1; Fig. 2B, C). The cancer cells exhibited a gene expression profile consistent with squamous epithelial differentiation. Upregulated genes included CDKN2A (p16), KRT6A, FSCN1, and SOX2, whereas CYP26A1, ACSS3, and ADH1C were among the most significantly downregulated genes. Expression of KRT5 and EPCAM was retained across all cancer samples, supporting the epithelial and basal characteristics of the tumor cells. In contrast, NKX2-1 (TTF1) expression was absent, further supporting squamous rather than adenocarcinoma differentiation (Fig. 2D). Pathway analysis revealed that the upregulated genes in cancer cells converged on proliferative programs. GSEA confirmed E2F targets, mTORC1 signaling, MYC targets, glycolysis, and G2M checkpoint as the most enriched Hallmark pathways (26,27) (Fig. 2G, Supplementary Table 6). To dissect the mechanistic basis of this proliferative switch, we examined the quiescence-maintaining pathways in turn.

**Figure 2.**
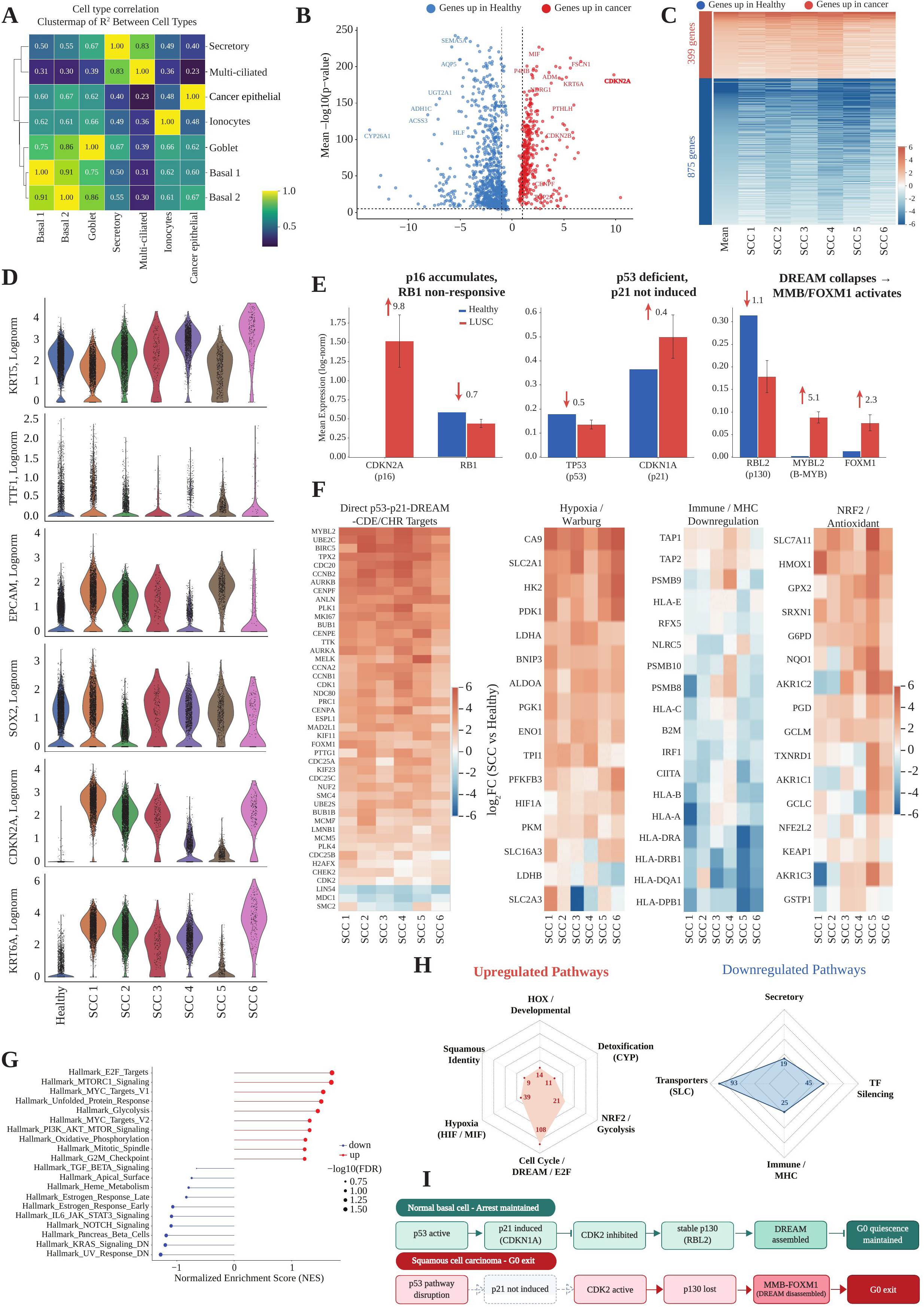
LUSC exhibits convergent disruption of the DREAM and RB1 quiescence axes. A, Clustermap of the coefficient of determination between cell types, showing hierarchical clustering of Basal 1, Basal 2, Goblet, Secretory, Multi-ciliated, Ionocyte, and Cancer epithelial transcriptomes. B, Volcano plot of differentially expressed genes between healthy and cancer epithelial cells, with the 875 genes upregulated in healthy tissue shown in blue and the 399 genes upregulated in cancer shown in red. C, Heatmap of the same 1274 differentially expressed genes across the six individual squamous cell carcinoma tumors, SCC1 through SCC6. D, Violin plots of KRT5, TTF1, EPCAM, SOX2, CDKN2A, and KRT6A expression in healthy tissue and in each of the six squamous cell carcinoma tumors. E, Bar plots comparing mean expression of CDKN2A and RB1, of TP53 and CDKN1A, and of RBL2, MYBL2, and FOXM1 between healthy and lung squamous cell carcinoma samples, with log2 fold change values indicated (log2FC +9.8 for CDKN2A and -0.7 for RB1; -0.5 for TP53 and +0.4 for CDKN1A; -1.1 for RBL2, +5.1 for MYBL2, and +2.3 for FOXM1), illustrating an accumulation of p16 without a corresponding decrease in proliferation. F, Heatmaps of log2 fold change in lung squamous cell carcinoma relative to healthy tissue across the six tumors for DREAM target genes, in addition to Hypoxia/Warburg effect genes, Immune/MHC related genes, and NRF2 antioxidant response genes. G, Lollipop plot of normalized enrichment scores for Hallmark gene sets significantly upregulated or downregulated in lung squamous cell carcinoma, with point size indicating statistical significance. H, Radar plots summarizing the number of genes contributing to upregulated pathway categories, including developmental (14 genes), detoxification (11 genes), NRF2 and glycolysis (21 genes), cell cycle and DREAM and E2F (108 genes), hypoxia (39 genes), and squamous identity (9 genes) programs, and to downregulated pathway categories, including secretory (19 genes), transcription factor silencing (45 genes), immune and MHC (25 genes), and transporter (93 genes) programs. I, Schematic summarizing the proposed mechanism by which G0 quiescence is established in normal basal cells via DREAM complex assembly, and G0 exit in lung squamous cell carcinoma driven by p53 pathway disruption and subsequent DREAM disassembly.

For the p53-DREAM axis, TP53 is mutated in approximately 80% of LUSC (4). Consistent with loss of function, its canonical target CDKN1A/p21 failed to be induced (Fig. 2E). Loss of RBL2/p130 (log2FC = -1.1) was accompanied by marked upregulation of MYBL2/B-MYB (log2FC = +5.1) and FOXM1 (log2FC = +2.3), indicating DREAM disassembly and switch to the proliferation-promoting MMB-FOXM1 complex (Fig. 2E, I). To quantify genome-wide DREAM disruption, we examined the 210 direct p53-p21-DREAM-CDE/CHR target genes defined by Fischer and Engeland (28). All 210 were detected in our data, of which 146 (70%) showed significant de-repression, while only 14 (7%) were downregulated (Fisher’s exact OR = 8.3, P = 3.3 x 10^-49^; Fig. 2F, Supplementary Fig. E6). This de-repression was consistent across all six tumors regardless of mutational background.

For the RB1 pathway, we observed a hallmark of functional RB1 inactivation: CDKN2A/p16 was among the most upregulated genes in the dataset (log2FC = +9.8) yet failed to arrest proliferation (Fig. 2E). This accumulation of p16 in the absence of cell cycle arrest indicates that the RB1 pathway is non-functional in these tumors (29). RB1 itself was reduced (log2FC = -0.7), also consistent with inactivation. In addition, E2F1/2/3 were significantly upregulated (log2FC approximately +2.2 (Supplementary Table 12)), confirming release from RB1 restraint.

Beyond cell cycle deregulation, cancer cells showed coordinated changes in metabolic and immune programs, including activation of hypoxia/Warburg metabolism CA9, SLC2A1, HK2, LDHA, PDK1 (Fig. 2F), downregulation of MHC class I/II and antigen-processing machinery TAP1/2, HLA-A/B/C, HLA-DRA/DRB1, B2M, IRF1 (Fig. 2F), and activation of the NRF2/antioxidant program SLC7A11, HMOX1, GPX2, SRXN1, G6PD, and NQO1 (Fig. 2F). These process-level changes converged into distinct functional modules (Fig. 2H), cell cycle/DREAM/E2F (108 genes), hypoxia/HIF/MIF (39 genes), NRF2/glycolysis (21 genes), squamous identity (9 genes), and HOX/developmental programs (14 genes) were upregulated, while secretory (19 genes), transporter (93 genes), detoxification, immune/MHC (25 genes), and Transcription factors (TF) (45 genes) were downregulated (Supplementary Fig. E8). These changes were consistent across all six tumors despite their distinct mutational landscapes, indicating convergent transcriptional rewiring rather than patient-specific alterations. The downregulation of MHC class I/II and antigen processing machinery may have implications for immune checkpoint therapy response in these tumors. TF silencing was among the most prominent downregulated programs (Fig. 2H), raising the question of which transcription factors normally maintain the balance between quiescence and proliferation in airway basal cells, and which are deregulated during squamous transformation. To address this, we next mapped the complete TF landscape of the healthy airway differentiation hierarchy.

### The basal compartment harbors four distinct states along a quiescence-to-activation gradient

To identify the transcription factors that maintain quiescence in airway basal cells, we computed diffusion pseudotime (30) across the healthy epithelium, starting from the TP63-high BSCs and terminating in multiciliated cells (Fig. 3A, C, Supplementary Fig. E9, Supplementary Table 7). The trajectory recapitulated the expected differentiation hierarchy. Lineage scoring confirmed that basal identity peaked during early pseudotime, followed sequentially by suprabasal, goblet, secretory, and ciliated programs (Fig. 3C). Sub-population analysis revealed heterogeneity within each major lineage (Fig. 3F). Closer examination of the basal compartment revealed additional states beyond the two populations identified by standard clustering (Fig. 1B). Pseudotime resolution uncovered four transcriptionally distinct states (Basal I-IV; Fig. 3D). Basal I cells expressed KRT15 and DST, while Basal III and IV showed progressively distinct profiles including ATF3, EGR1, CLDN4, and CD55 (Fig. 3E) (Supplementary Table 8, 9). Pearson correlation confirmed that these four populations form a differentiation gradient: highly correlated with each other (r = 0.95-0.98) but progressively divergent from multiciliated cells (r = 0.70-0.73; Supplementary Fig. E11). The high inter-state correlations indicate a graded continuum rather than four discrete cell types, consistent with the transitional-state model derived from single-cell cloning of human bronchial basal cells (14). Lineage scoring across the basal region showed distinct peaks corresponding to each population (Fig. 3I). These programs were also conserved between young and aged donors (Fig. 3J, Supplementary Fig. E12), indicating that the substructure reflects stable biology rather than age-related drift.

**Figure 3.**
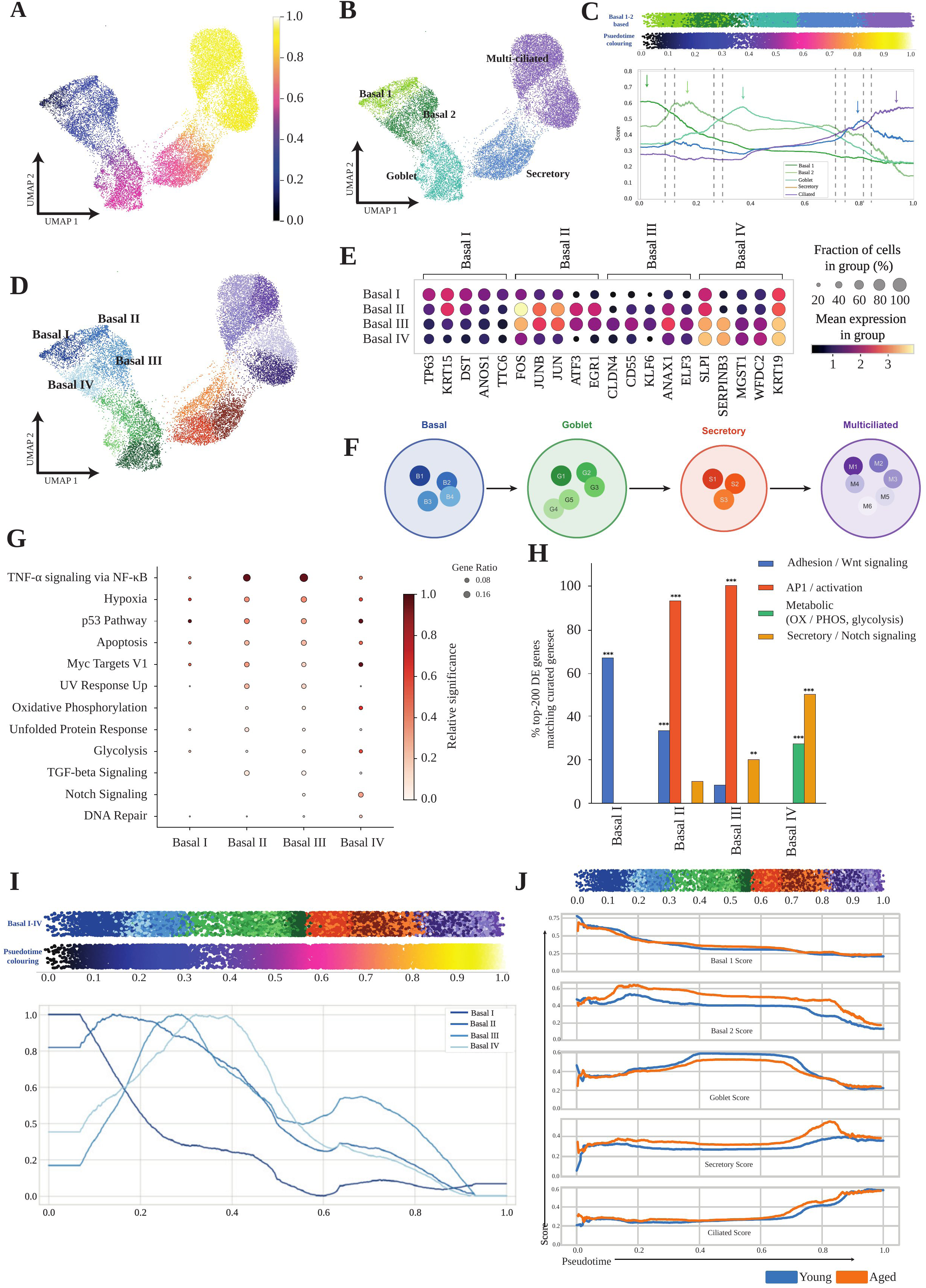
The basal compartment harbors four distinct states along a quiescence to activation gradient. A, UMAP plot of the airway epithelial differentiation trajectory colored by pseudotime, with the starting point shown in dark blue (0.0). B, The same UMAP embedding colored by the major annotated cell types, Basal 1, Basal 2, Goblet, Secretory, and Multi-ciliated. C, Lineage identity scores for the basal, suprabasal, goblet, secretory, and ciliated programs plotted along pseudotime, with the pseudotime rank and cell type identity of each cell shown as colored strips above the plot and dashed vertical lines and arrows marking inflection points in each lineage score. D, UMAP plot of the basal compartment subdivided into four subclusters, Basal I, Basal II, Basal III, and Basal IV. E, Dot plot of marker genes defining Basal I through Basal IV, grouped by the top differentially expressed genes for each subcluster. F, Schematic of the proposed basal to goblet to secretory to multiciliated differentiation lineage, depicting individual basal, goblet, secretory, and multiciliated subpopulations as connected nodes. G, Dot plot of Hallmark pathway enrichment across Basal I through Basal IV. H, Bar chart of the percentage of the top two hundred differentially expressed genes for each basal subcluster matching curated adhesion/Wnt signaling, AP-1/activation, metabolic (oxidative phosphorylation/glycolysis), and secretory/Notch signaling gene sets, with significance levels indicated. I, Basal I through Basal IV identity scores plotted along pseudotime, with the pseudotime rank and cell type identity strips shown above. J, Basal, suprabasal, goblet, secretory, and ciliated lineage scores plotted along pseudotime and compared between young and aged donors, with the pseudotime rank and cell type identity strips shown above.

Marker gene expression and Hallmark GSEA and functional gene set enrichment revealed distinct biological roles for each state (Fig. 3G) (Supplementary Table 9). Basal I showed a signature of Wnt signaling and cell adhesion consistent with stem cell quiescence (31,32), Basal II showed early activation of AP-1 genes (Activated), Basal III displayed peak activation with maximal AP-1 gene activity and metabolic programs (Hyperactivated), and Basal IV was enriched for secretory and Notch signaling (Primed) (Fig. 3H). This resolution demonstrates that basal cell activation follows a continuous trajectory, with quiescence confined to the subpopulation at the top of the pseudotime hierarchy. Basal I occupies the earliest pseudotime position and lacks the AP-1, metabolic, and secretory or Notch signaling signatures that define Basal II through IV. This quiescent profile, defined by Wnt signaling and cell adhesion, is consistent with the stem cell state described for basal cells in the airway epithelium (8), raising the question of which transcription factor establishes and maintains this quiescent identity.

### HLF is a quiescence-associated transcription factor in the basal stem cell compartment, silenced in lung squamous cell carcinoma

Having resolved four basal states with distinct biological identities, we asked which transcription factor programs are retained or lost in cancer. We grouped TFs by their pseudotime program and compared expression across Quiescent, Activated, Hyperactivated, Primed, and Cancer epithelial cells (Fig. 4K). Of 441 total and 173 basal-specific TFs mapped along the differentiation trajectory (Supplementary Fig. E14 and E15), cancer cells shared maximum similarity with the Hyperactivated program including MYC, FOS, JUN, FOSB, and EGR1, while 18 of 27 Basal I-associated TFs were downregulated, several known to inhibit cell cycle, indicating systematic silencing of the quiescence program. To understand what these silenced TFs normally do, we examined their expression along the pseudotime in healthy tissue (Fig. 4B-E). Known basal regulators occupied expected positions, TP63, the master regulator of basal cell identity, was expressed from the earliest pseudotime through Basal I-III (Fig. 4B), consistent with its established role in airway stem cell maintenance (33). Immunofluorescence confirmed TP63 and KRT5 co-localization in basal cells (Fig. 4F). Proliferative markers E2F1, TOP2A, and MKI67 were confined to Basal II-IV but absent from Basal I (Fig. 4G), confirming the quiescent state of the most primitive population and providing a clear rationale for identifying the TFs that distinguish Basal I from its neighbors.

**Figure 4.**
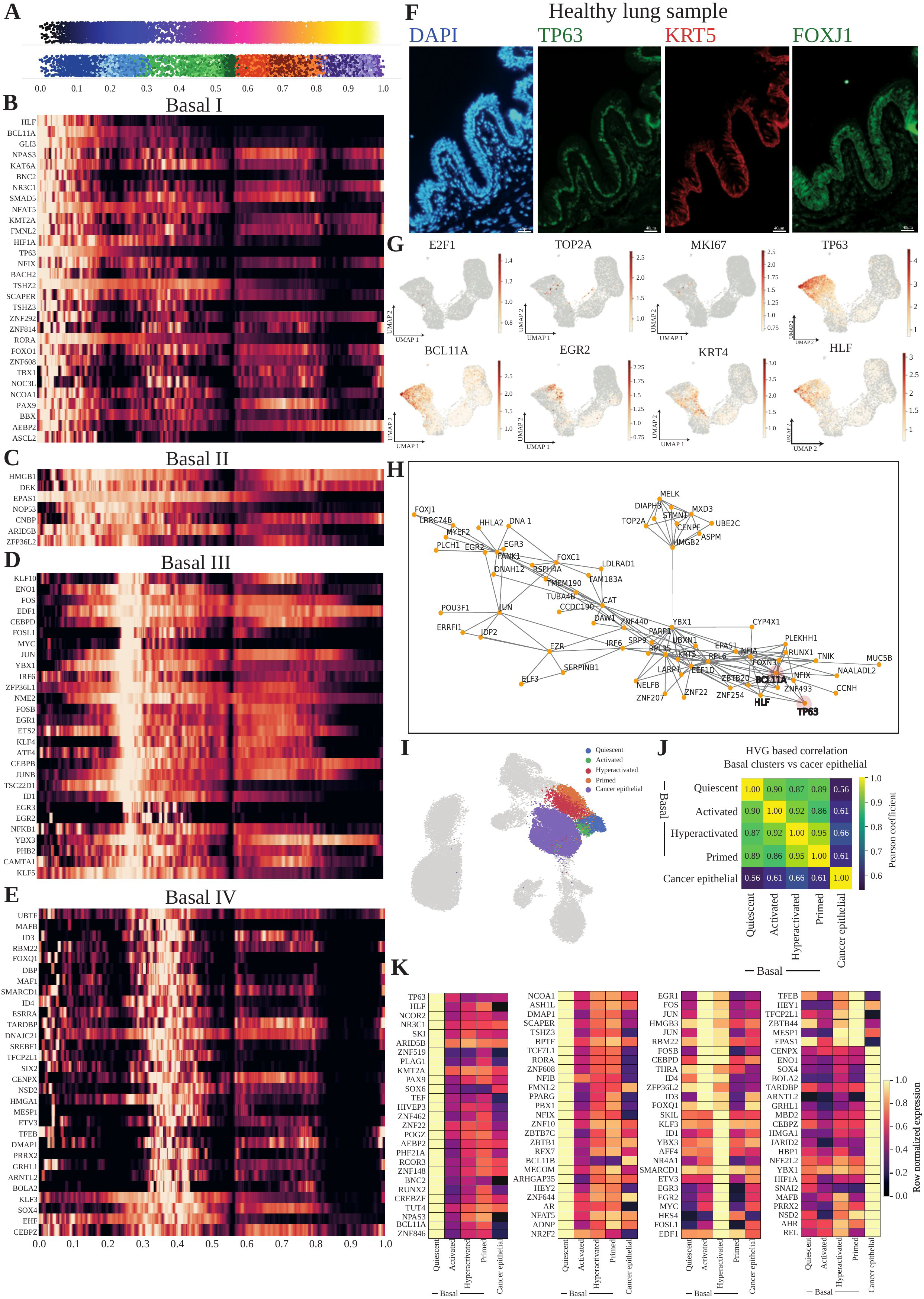
HLF is a quiescence-associated transcription factor in the primitive basal compartment, silenced in LUSC. A, Pseudotime rank and cell type identity strips shown as a reference for the heatmaps in B through E. B, Heatmap of transcription factor expression along pseudotime in the Basal 1 subcluster, including HLF, BCL11A, GLI3, and related factors. C, Heatmap of transcription factor expression along pseudotime in the Basal 2 subcluster. D, Heatmap of transcription factor expression along pseudotime in the Basal 3 subcluster. E, Heatmap of transcription factor expression along pseudotime in the Basal 4 subcluster. F, Immunofluorescence images of a healthy lung tissue section stained for DAPI, TP63, KRT5, and FOXJ1. G, UMAP feature plots of E2F1, TOP2A, MKI67, TP63, BCL11A, EGR2, KRT4, and HLF expression in the basal compartment. H, Gene regulatory correlation network of transcription factors and correlated genes in the basal compartment, with HLF, TP63, and BCL11A highlighted as interconnected hub nodes. I, UMAP plot of the basal and cancer epithelial compartment colored by five substates, Quiescent, Activated, Hyperactivated, Primed, and Cancer epithelial. J, Heatmap of pairwise Pearson correlation coefficients, calculated from highly variable genes, between the Quiescent, Activated, Hyperactivated, and Primed basal substates and Cancer epithelial cells, with coefficients ranging from 0.86 to 0.95 among the basal substates and from 0.56 to 0.66 between the basal substates and Cancer epithelial cells. K, Heatmaps of row normalized transcription factor expression across the Quiescent, Activated, Hyperactivated, Primed, and Cancer epithelial groups, including HLF, TP63, and related transcription factors.

The Basal I program comprised 89 TFs expressed at the earliest pseudotime (Fig. 4B, Supplementary table 8), including HLF, BCL11A, GLI3, NFAT5, KMT2A, BNC2, SMAD5, NFIA, and NFIB, most of which have been previously linked to airway basal cell quiescence. The Basal II program contained 7 TFs enriched for chromatin modulators (Fig. 4C), including HMGB1, EHF, and ARID5B. The Basal III program was dominated by immediate-early response factors (MYC, FOS, JUN, FOSB, EGR1); notably, MYC expression spanned both Basal II and III. The Basal IV program included TFs linked to differentiation commitment (Fig. 4E), including DBP, ESRRA, and SOX4.

Gene co-expression network analysis (SCENIC) (34) revealed the regulatory architecture of the quiescent basal state. HLF, TP63, and BCL11A occupied highly connected positions in the network, while also having shared regulatory targets (Fig. 4H). Correlation analysis confirmed that cancer cells were most similar to the “hyperactivated” state and least similar to the “quiescent” state (Fig. 4I,J). UMAP projection confirmed that HLF expression was restricted to the Basal I compartment, overlapping with TP63 in healthy tissue but absent from cancer epithelial cells (Fig. 4G, bottom). Among all Basal I TFs, HLF showed the most consistent silencing across tumors. In addition, its family member TEF (log2FC = -2.9, 5/6) and the circadian bHLH factor BHLHE41 (log2FC = -4.3, 6/6 patients) (Supplementary Table 5) were also downregulated, suggesting coordinated silencing of circadian and quiescence-associated factors during LUSC development. Together, these data provide a comprehensive transcription factor landscape of the human airway basal compartment and, through the full epithelial hierarchy presented in the Supplementary data, a resource that can guide future studies of airway biology and disease. Within this landscape, HLF emerged as the most consistently silenced quiescence factor across all six tumors, identifying it as a candidate for functional validation.

### HLF reinstates quiescence in basal stem cells through selective DREAM target suppression

Having identified HLF as a candidate quiescence factor silenced in LUSC, we tested whether its re-expression could reinstate quiescence. Consistent with a role in quiescence, HLF was expressed in freshly isolated BSCs but lost when BSCs proliferated in culture (Fig. 5A). If HLF maintains quiescence, forced re-expression should arrest proliferating cells. To test this, we overexpressed HLF using a lentiviral vector, kindly provided by the Bryder lab, (24) (Fig. 5B) in primary BSCs and in a BSC line with full differentiation capacity BCi-NS1 (Supplementary Fig. E16) (35). HLF-expressing cells showed progressive loss of ZsGreen+ frequency over eight days (Fig. 5C, D, Supplementary Fig. E17). Cell-cycle analysis at day 4 revealed that HLF induced G0 quiescence: the fraction of cells in G0 increased from 0.73% to 38.2%, while G1 cells decreased correspondingly (68.6% to 27.6%; Fig. 5E-G).

**Figure 5.**
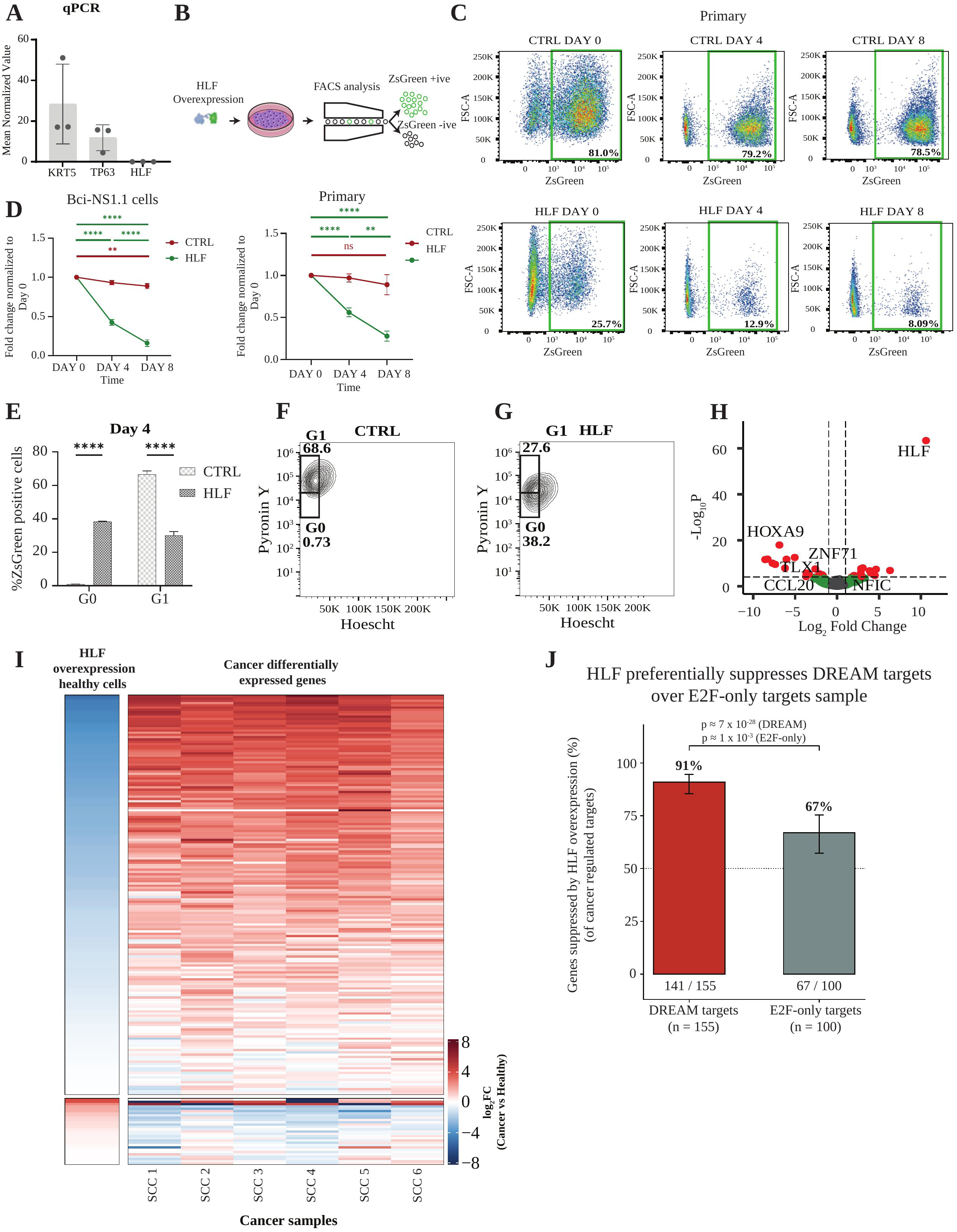
HLF reinstates quiescence in basal stem cells through selective DREAM target suppression. A, Bar plot of mean normalized KRT5, TP63, and HLF expression confirming basal cell identity and low baseline HLF expression in Primary cells prior to transduction. B, Schematic of the lentiviral HLF overexpression and ZsGreen fluorescent reporter workflow, from transduction through fluorescence-activated cell sorting into ZsGreen-positive and ZsGreen-negative populations. C, Representative flow cytometry plots of ZsGreen fluorescence in control and HLF-transduced Primary cells at Day 0, Day 4, and Day 8, with 81.0%, 79.2%, and 78.5% ZsGreen-positive cells in control and 25.7%, 12.9%, and 8.09% ZsGreen-positive cells in the HLF condition at the same time points. D, Growth curves showing fold change in cell number normalized to Day 0 for control and HLF-overexpressing BCi-NS1.1 cells and for control and HLF-overexpressing Primary cells, with statistical comparisons across time points. E, Bar plot of the percentage of ZsGreen-positive cells in the G0 and G1 cell cycle phases at Day 4 for control and HLF-overexpressing cells. F, Representative Pyronin Y and Hoechst flow cytometry plot identifying G0 (0.73%) and G1 (68.6%) populations in control cells. G, Representative Pyronin Y and Hoechst flow cytometry plot identifying G0 (38.2%) and G1 (27.6%) populations in HLF-overexpressing cells, showing an increased G0 fraction relative to control. H, Volcano plot of genes differentially expressed between HLF-overexpressing and control cells by bulk RNA sequencing, with HLF, HOXA9, ZNF71, TLX1, CCL20, and NFIC among the most significantly differentially expressed genes. I, Heatmap comparing the transcriptional signature of HLF overexpression in healthy cells with the cancer differentially expressed gene signature across the six squamous cell carcinoma tumors, SCC1 through SCC6, showing that genes downregulated by HLF overexpression correspond largely to genes upregulated in cancer and vice versa. J, Bar plot showing that HLF overexpression preferentially suppresses DREAM complex target genes compared with E2F-only target genes, with 91% (141 of 155) of DREAM target genes suppressed compared with 67% (67 of 100) of E2F-only target genes, and statistical significance indicated (p ≈ 7 × 10⁻²⁸ for DREAM targets and p ≈ 1 × 10⁻³ for E2F-only targets). Error bars represent 95% binomial confidence intervals.

Bulk RNA-seq of HLF-overexpressing BCi-NS1 cells at 48 hours confirmed downregulation of cell cycle genes and known HLF regulated pathways (Supplementary Fig. E18, Supplementary Table 10). We also identified HOXA9 as the most suppressed gene, alongside TLX1, ZNF71, and CCL20 (Fig. 5H). To directly test whether HLF suppresses the DREAM proliferative program, we compared genes suppressed by HLF with genes de-repressed in cancer. The dual heatmap (Fig. 5I) demonstrated a clear inverse relationship: genes upregulated in cancer (right panel, red) were suppressed by HLF overexpression in healthy cells (left panel, blue), while genes downregulated in cancer showed the opposite pattern. Of 181 detectable DREAM-CDE/CHR targets in the RNA-seq, 155 (86%) were upregulated in LUSC. Of these 155, 141 (91%) were suppressed upon HLF overexpression (binomial P = 7.0 x 10-28; median FC = 0.79; Fig. 5I). To assess pathway specificity, we compared this with E2F-only targets (MSigDB Hallmark, excluding shared genes): only 67% of upregulated E2F-only targets were suppressed by HLF (median FC = 0.92). This preferential suppression of DREAM targets (141 of 155 DREAM-CDE/CHR targets suppressed, 91%, versus 67 of 100 E2F-only targets, 67%; Fig. 5J), indicating that HLF acts through the DREAM/p130 pathway rather than the RB1/E2F axis.

### HLF suppresses proliferation in LUSC and predicts patient survival

Finally, to test whether HLF can reinstate quiescence in LUSC cells, we selected two TP53-mutant cell lines, HCC95 and HCC1588 (Korean Cell Line Bank), neither of which expresses appreciable HLF (Fig. 6A, B). Lentiviral transduction achieved re-expression of HLF with transduction efficiencies of 32.3% and 58.5% respectively (Fig. 6C), and proliferation assays demonstrated significant, sustained growth suppression in both cell lines over eight days (Fig. 6D, H, Supplementary Fig. E19). Despite substantial genetic divergence between these lines (Supplementary Fig. E20), both responded similarly, consistent with a mechanism independent of specific mutational context.

**Figure 6.**
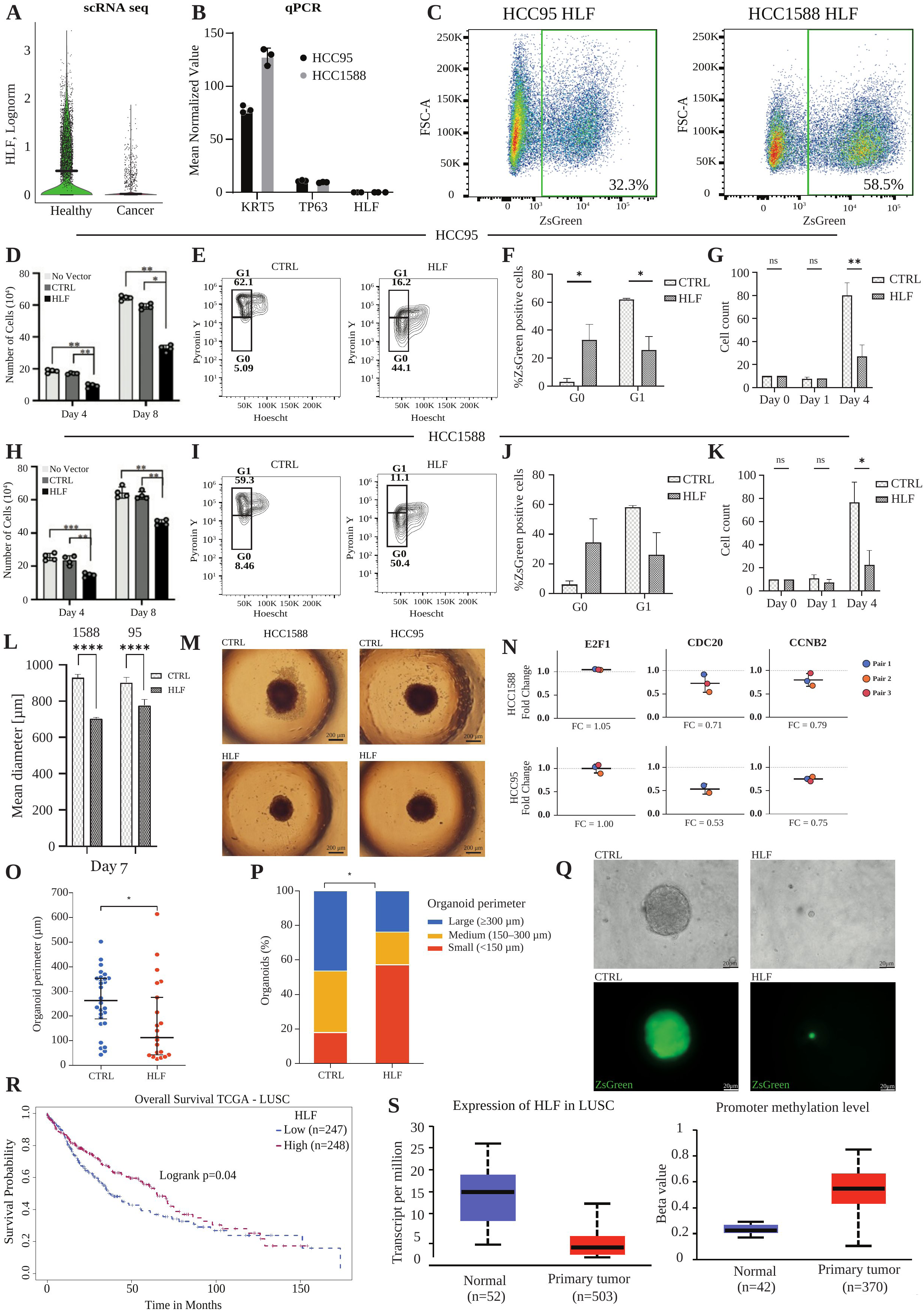
HLF suppresses proliferation in LUSC and predicts patient survival. A, Violin plot of HLF expression by single cell RNA sequencing in healthy and cancer epithelial cells, showing markedly reduced HLF expression in cancer. B, Bar plot of mean normalized KRT5, TP63, and HLF expression in the HCC95 and HCC1588 lung squamous cell carcinoma cell lines prior to transduction. C, Representative flow cytometry plots of ZsGreen fluorescence following HLF lentiviral transduction, showing 32.3% ZsGreen-positive cells in HCC95 and 58.5% in HCC1588 cells. D, Bar plot of cell numbers at Day 4 and Day 8 in HCC95 cells for no vector, control, and HLF-overexpressing conditions. E, Pyronin Y and Hoechst flow cytometry plots of control (G0 5.09%, G1 62.1%) and HLF-overexpressing (G0 44.1%, G1 16.2%) HCC95 cells. F, Bar plot of the percentage of ZsGreen-positive cells in G0 and G1 in control and HLF-overexpressing HCC95 cells. G, Bar plot of cell counts at Day 0, Day 1, and Day 4 in control and HLF-overexpressing HCC95 cells. H, Bar plot of cell number at Day 4 and Day 8 in HCC1588 cells for no vector, control, and HLF-overexpressing conditions. I, Pyronin Y and Hoechst flow cytometry plots of control (G0 8.46%, G1 59.3%) and HLF-overexpressing (G0 50.4%, G1 11.1%) HCC1588 cells. J, Bar plot of the percentage of ZsGreen-positive cells in G0 and G1 in control and HLF-overexpressing HCC1588 cells. K, Bar plot of cell counts at Day 0, Day 1, and Day 4 in control and HLF-overexpressing HCC1588 cells. L, Bar plot of mean spheroid diameter at Day 7 in control and HLF-overexpressing HCC95 and HCC1588 cells. M, Representative brightfield images of control and HLF-overexpressing spheroids from HCC1588 and HCC95 cells. N, Quantitative PCR fold change of E2F1, CDC20, and CCNB2 in HLF-overexpressing relative to control HCC1588 (FC = 1.05, 0.71, and 0.79, respectively) and HCC95 (FC = 1.00, 0.53, and 0.75, respectively) cells across three paired experiments. O, Violin plot of spheroid perimeter in control and HLF-overexpressing conditions. P, Stacked bar plot of the fraction of spheroids falling into small (≤150 μm), intermediate (100–300 μm), and large (>300 μm) perimeter categories in control and HLF-overexpressing conditions. Q, Brightfield and TP63 immunofluorescence images of representative control and HLF-overexpressing spheroids. R, Kaplan-Meier overall survival curve from The Cancer Genome Atlas lung squamous cell carcinoma cohort comparing patients with low (n=247) and high (n=248) HLF expression, with a logrank p value of 0.04. S, Box plots of HLF transcript abundance (normal n=52, primary tumor n=503) and HLF promoter methylation beta value (normal n=42, primary tumor n=370) in normal and primary tumor samples from The Cancer Genome Atlas lung squamous cell carcinoma cohort.

Cell-cycle profiling revealed that HLF drove both lines into G0 quiescence: G0 fractions rose from 5.09% to 44.1% (HCC95) and from 8.46% to 50.4% (HCC1588), with corresponding reductions in G1 (62.1% to 16.2% and 59.3% to 11.1%, respectively; Fig. 6E, F, I, J, Supplementary Fig. E21). To isolate the effect of HLF from non-transduced neighbors, we sorted transduced cells and plated them at clonal density. HLF-expressing cells remained viable but failed to form colonies over four days (Fig. 6G, K), confirming cell-autonomous growth arrest rather than toxicity. Spheroid growth was also significantly reduced in both lines, with mean spheroid diameter decreasing over seven days (Fig. 6L, M). To confirm pathway specificity in LUSC cells, we measured E2F1 (RB1/E2F pathway readout) and two direct DREAM-CDE/CHR targets (CDC20, CCNB2) by qRT-PCR 48 hours post-transduction of HLF and cotrol vector (Fig. 6N). E2F1 was unchanged in both lines (FC = 1.05 and 1.00), while CDC20 and CCNB2 showed consistent suppression (FC = 0.53-0.79), indicating early and selective suppression of DREAM targets rather than the RB1/E2F program. This early timepoint captures direct transcriptional effects before secondary proliferation changes confound the analysis.

Critically, HLF also suppressed growth in patient-derived LUSC organoids, reducing organoid size and shifting the distribution toward smaller organoids (Fig. 6O, P). Transduced organoids retained TP63 expression (Fig. 6Q, Supplementary Fig. E22), demonstrating that HLF enforces quiescence without compromising basal identity. Finally, to assess clinical relevance, we interrogated TCGA expression and methylation data. LUSC tumors showed markedly reduced HLF transcript levels (∼15 to ∼5 TPM, n = 503 vs 52 normal) and significantly elevated HLF promoter methylation (beta value ∼0.22 in normal tissue vs ∼0.60 in primary tumors, n = 370 vs 42; Fig. 6S), identifying promoter hypermethylation as associated with HLF silencing. Patients with low HLF expression exhibited significantly worse overall survival (n = 247 low vs n = 248 high; log-rank P = 0.04; Fig. 6R). These findings independently validate HLF as a clinically relevant tumor suppressor whose silencing mechanism is potentially amenable to pharmacological intervention.

Together, these data demonstrate that HLF reinstates quiescence across multiple LUSC model systems, acts through DREAM target suppression rather than the RB1/E2F axis, and carries prognostic significance in an independent patient cohort.

## Discussion

This study identifies HLF as a quiescence-governing transcription factor silenced by promoter hypermethylation in LUSC whose re-expression is sufficient to reinstate quiescence in cancer cells with TP53 loss. By integrating single-cell atlases of the healthy airway and LUSC with functional validation across primary BSCs, LUSC cell lines, and patient-derived organoids, we demonstrate that HLF maintains quiescence in the earliest basal stem cell population, and that its forced re-expression selectively suppresses DREAM complex targets to drive G0 arrest.

LUSC is dominated by loss-of-function alterations, most prominently TP53 mutation and RB pathway disruption (4), rather than the targetable gain-of-function drivers found in lung adenocarcinoma (4). Our data demonstrate the cellular basis of this proliferative state. Proliferation is driven by two converging checkpoint losses. DREAM complex disruption, evidenced by de-repression of 146 of 210 direct CDE/CHR target genes, eliminates G0 maintenance, while RB1 inactivation, consistent with marked p16 accumulation that fails to arrest proliferation, eliminates the G1/S checkpoint. These two pathways control distinct gene sets at distinct checkpoints: DREAM represses G2/M genes to maintain G0, while RB1 sequesters E2F1/2/3 to block G1/S entry (36)(36). Both must function for quiescence; neither does in the tumors we examined. This provides direct cellular evidence for the failure of CDK4/6 inhibitors in LUSC(21), which require functional RB1 to mediate their effects (37).

Our resolution of four transcriptionally distinct states within the basal compartment provides new insight into the origin of the LUSC proliferative state. Rather than a simple binary between quiescent basal cells and proliferative suprabasal cells, we find a gradient from the most primitive quiescent state (Basal I, expressing HLF, BCL11A, GLI3) through transitional states to an activated/proliferative state (Basal III, expressing MYC, FOS, JUN, FOSB, EGR1). Cancer cells most closely resemble this activated Basal III state, suggesting that transformation locks cells in a transit-amplifying-like program while silencing the Basal I quiescence module. Slow cycling in LUSC can arise through more than one program, a KRT13+ hillock-like population driven by KLF4 has recently been described across squamous models and linked to platinum resistance (38). That state is defined by barrier identity and drug tolerance, whereas the Basal I module described here is the quiescence program of the normal basal hierarchy, silenced in tumors rather than adopted by them.

The identification of HLF as a Basal I-specific quiescence factor provides a principled basis for therapeutic TF selection. Rather than screening arbitrary candidates, our pseudotime and network analysis placed HLF at the core of the quiescent basal regulatory network, alongside TP63 and BCL11A. The preferential suppression of DREAM targets (91%) over E2F-only targets (67%) links HLF to DREAM-mediated gene repression. The 210 Fischer DREAM targets were originally defined as p53-dependent (20,28), yet HLF suppressed DREAM targets CDC20 and CCNB2 also in the TP53-mutant lines HCC95 and HCC1588 while leaving E2F1 unchanged (Fig. 6N), indicating that HLF engages DREAM target repression independently of p53, though whether HLF directly promotes DREAM complex assembly or acts through an intermediate pathway remains to be determined.

Because HLF loss is associated with promoter methylation rather than genetic deletion, it remains structurally intact and could potentially be reactivated by DNA methyltransferase inhibitors already in clinical use for hematological malignancies (39,40). Monotherapy with these agents has so far shown modest activity in solid tumors, so combination regimens or more selective reactivation strategies are likely to be required (41). Whether demethylating agents can selectively restore HLF expression in LUSC warrants investigation.

The coordinated loss of multiple circadian and quiescence-associated factors (HLF, TEF, BHLHE41) alongside upregulation of the polycomb repressor EZH2 suggests active epigenetic remodeling of quiescence-regulatory programs during LUSC development. HOXA10 was among the most upregulated genes in our LUSC dataset and was previously shown to directly regulate HLF expression (40). Here we find the reciprocal arm, HLF overexpression suppresses HOXA9 as its most downregulated gene. Together, these observations suggest a regulatory loop in which HOX gene activation and HLF loss may be mutually reinforcing, though direct evidence for this circuit remains to be established.

HLF as a tumor suppressor is consistent with emerging evidence, including suppression of metastasis in NSCLC (42). However, HLF function is context-dependent, in hepatocellular carcinoma (43) and intrahepatic cholangiocarcinoma (44) it has been reported to act as an oncogene, while in AML it maintains leukemic stem cell quiescence (45). This duality is consistent with the expected behavior of a quiescence factor, tumor-suppressive in epithelial cancers where quiescence restrains proliferation, but potentially co-opted in hematopoietic malignancies where quiescence protects the leukemic stem cell pool. HLF has also been shown to direct tissue residency of proinflammatory memory CD4+ T cells (46), indicating that it sets cell state across lineages rather than through a single effector program.

Published studies of DREAM manipulation have focused on disrupting the complex to release proliferation, in lung tumorigenesis (47), alveolar regeneration (48), and pancreatic beta cells (49). Our findings demonstrate the reverse, a tissue-specific transcription factor can restore DREAM-target repression and impose G0 arrest. In summary, HLF is a quiescence factor and its reactivation suppresses DREAM target gene expression and reinstates G0 arrest regardless of mutational context. By restoring a transcriptional program rather than targeting individual genetic lesions, HLF reactivation offers a potential strategy for a cancer that has resisted genomics-driven drug development.

## Methods

For detailed descriptions, see Supplementary Methods and supplementary table 11 for reagents.

### Ethical approval

Primary healthy and tumor cells were obtained from bronchoscopy or explants as previously described (50). Bronchoscopies were performed under local anaesthetics in accordance with clinical routines. All patient material was transported, on ice in cancer cell media, from the clinic within a few hours of the surgery. All donors gave informed consent, and ethical approval for the study was obtained from the Swedish Ethical Review Authority (ethical permit numbers 2014/32, 2017/180, and 2017/519).

### Cell Culture

In Vitro cultures of basal cells and cancer cells were grown in PneumaCult-Ex media and DMEM + 10% FBS media respectively (see supplementery table 11). Both media were supplemented with penicillin/streptomycin (Fisher Scientific). At 70% confluence, cells were passaged (1:4) and replated. ALI cultures were set up in accordance with the protocol from STEMCELL technologies using the PneumaCult-ALI basal medium to differentiate the cells.

### Single-cell RNA sequencing

Single-cell RNA sequencing (scRNAseq) was performed by the Single-Cell Genomics Platform at the Center for Translational Genomics at Lund University. Single cell fractions of our fresh frozen airway epithelial cells and tumor cells were generated after the cells were thawed and then processed to remove debris, doublets and dead cells. The tumor cells were additionally processed to remove white blood cells. scRNAseq libraries were constructed using the Chromium Single Cell Gene Expression 3’ v3 Reagent Kit according to the manufacturer’s instructions. These libraries were then sequenced on a NovaSeq 6000 or a Nextseq 500 (Illumina).

### Processing of single cell RNA seq data

Primary processing was performed using the Cell Ranger pipeline (v6.1, 10x Genomics). Secondary pre-processing and downstream analysis were performed within the Scanpy ecosystem. Cells with fewer than 1,000 detected transcripts (unique molecular identifiers, UMIs) or with >12% of reads mapping to mitochondrial genes were excluded. Gene expression values were log-transformed and normalized to the median expression across the dataset. Uniform Manifold Approximation and Projection (UMAP) was applied for two-dimensional visualization. To assess disruption of the DREAM and E2F dependent programs we utilized the curated target genes defined by Fischer and Engeland (28).

Differentially expressed genes (DEGs) were identified using a two-sided Wilcoxon rank-sum test to assess both cell type-specific differences (each cell type versus all others) and age-related differences (aged versus young samples). Gene Set Enrichment Analysis (GSEA) was conducted using 1000 gene set permutations under the same settings. Diffusion pseudotime was computed using a root cell defined by the highest expression of TP63 within the basal cell cluster. All data can be found under the Gene Expression Omnibus accession number: GSE295552.

### Immunohistochemistry

Formalin fixed paraffin embedded primary tissue samples were sectioned (5µm) and adhered to glass slides. The tissue sections were deparaffinized and subsequently immunostained with antibodies from Supplementary Table 11 and imaged using VS120 Virtual Slide microscope (Olympus) and processed using the OlyVIA software V3.2.1. Slides were also imaged on the IX73 Inverted Microscope (Olympus) using the cellSense standard software. ALI sections were formalin fixed and paraffin embedded. In short, ALI membranes were coated in liquid paraffin at 55°C for 25 min, flattened into wax chips, and embedded perpendicular to the blade. Sections of 8 µm were processed for analysis as above.

### Lentiviral induction and FACS analysis

To induce hepatic leukemia factor (HLF) overexpression, we used the lentiviral vector pHAGE2-Hlf/Hlf-IRES-ZsGreen (kindly provided by Bryder lab)(24). The virus was titrated by qPCR and applied at 30,000 genome copies per cell. To test proliferation in the presence of this vector, ZsGreen positive cells were analyzed through fluorescence activated cell sorting (BD AriaIII using a 85-μm nozzle). All proliferation experiments were performed using cytometers (Becton Dickinson) and analyzed by Flow Jo software (Tree Star, version 9.5.1 or 10.0.2). ZsGreen fluorescence was detected using the FITC channel.

### Cell cycle FACS analysis

Cell cycle state was further assessed by Pyronin Y and Hoechst 33342 co-staining, distinguishing a G0 population from a G1 population by flow cytometry. Bulk RNA sequencing was performed on BCi-NS1 cells 48 hours after HLF or control transduction to identify genes suppressed by HLF overexpression, which were compared against genes dysregulated in cancer. Quantitative RT-PCR (C1000 Touch Thermal Cycler (Bio-Rad)) was used to measure E2F1, CDC20, and CCNB2 transcript levels 48 hours after HLF transduction in HCC95 and HCC1588 cells. The data were normalized, and fold change was calculated as the ΔΔCT method.

### Spheroid culture

Cell line derived spheroids were cultured and seeded at 1,000 cells/well in 200 µL medium in 96-well low-adherence plates using a suspension culture protocol (see Supplementary Methods). The spheroids were imaged using an Olympus IX70 inverted microscope. Brightfield images were used to compute the mean diameter using Image J.

### Primary lung cancer organoid

Fresh lung tumor tissue was minced into 1–2 mm pieces followed by enzymatic digestion at 37 °C. The suspension was then strained (100 µm) and cells were embedded in Matrigel (65–70% Matrigel, 30–35% medium) as 50–60 µl domes in pre-warmed 24-well plates. Once the domes had set, they were overlaid with complete organoid medium containing Y-27632 for the first 3–5 days. Once established, organoids were dissociated and transduced with HLF or control vector in the presence of polybrene. The cell-virus mixture was sedimented by centrifugation (400g for 30 min) before being resuspended and re-embedded in Matrigel, to assess effects on organoid size distribution and TP63 expression. For the analysis, organoids were fixed in 2% paraformaldehyde overnight at 4°C. OCT compound (500 µL) was added per well, and organoid-containing OCT was scraped and transferred to 15 × 15 mm cryomolds. Molds were centrifuged at 350 × g for 5 min to settle organoids into a single focal plane and stored at −80°C. Cryosections of 5 µm were collected on adherent slides and imaged on an Olympus IX70 inverted fluorescence microscope.

### TCGA

HLF expression, HLF promoter methylation, and overall survival were analyzed using the TCGA lung squamous cell carcinoma cohort. https://ualcan.path.uab.edu/analysis.html

### Statistics

Differential expression across all single-cell RNA-sequencing analyses was assessed using a two-sided Wilcoxon rank-sum test. Cell-cycle flow cytometry experiments were analyzed by two-way ANOVA with Tukey post-hoc comparisons using the GraphPad Prism 10 software. Gene-set enrichment among DREAM/E2F targets was assessed with one-sided Fisher’s exact and binomial tests, and overall survival was compared using a log-rank test.

## Data Availability

Single-cell RNA-seq data have been deposited in the Gene Expression Omnibus under accession GSE295552. All other data supporting the findings of this study are available in the article and its supplementary information, or from the corresponding author upon reasonable request.

## Acknowledgments

We thank the Center for Translational Genomics (CTG) at Lund University for sequencing services and the StemTherapy FACS Core Facility for flow cytometry support.

## Author Contributions

Prabhala and M. Magnusson conceived and designed experiments; P. Prabhala, S. Freiman, J. Stevrell, J. Sharma, N. Gvazava, F. Rosengren, C. Vazquez Echegaray, E. Arbajian, S. Wijk, S. Lang, R. Cattani performed experiments; K. Kanzenbach, E. Tufvesson, M. Planck, and L. Bjermer provided samples; D. Bryder provided the lentiviral vector; H. Brunnström performed pathological assessment; P. Prabhala, S. Freiman, J. Sharma, J. Stevrell, N. Gvazava, S. Lang, M. Magnusson analyzed data; P. Prabhala, S. Freiman, S. Soneji, and S. Lang performed bioinformatic analysis. P. Prabhala, S. Freiman, J. Sharma, J. Stevrell, S. Lang, N. Gvazava, R. Cattani, D. Bryder, E. Arbajian, H. Brunnström, M. Planck, J. Flygare, J. Wigen, D. Wagner, G. Westergren Thorsson, M. Magnusson interpreted results. P. Prabhala, S. Freiman, J. Sharma, J. Stevrell, M. Magnusson prepared figures; P. Prabhala and M. Magnusson drafted the manuscript. All authors have read, reviewed, and approved the final manuscript.

