## Supplementary table legends for "HLF suppresses DREAM target genes and reinstates quiescence in lung squamous cell carcinoma"

Supplementary Table 1.

Clinical, demographic and molecular characteristics of the six LUSC patients and ten healthy donors. Sample ID, year of birth, age, age classification, sex and smoking status for all 16 subjects, plus, for the six tumors, pathological stage, fusion/amplification testing (EGFR, BRAF, ALK, ROS1, KRAS, RET), mutation testing (p53 in all six; FGFR3 additionally in Cancer 3), and immunohistochemical staining (CK5, p40, TTF-1).

Supplementary Table 2.

Pulmonary function and peripheral blood characteristics of the ten healthy donors. Year of birth, age, age classification, year of biopsy and sex for each donor, together with spirometry (FEV1, FEV1 % predicted, FEV1/FVC, DLCO % predicted) and peripheral blood differential counts (leukocytes, neutrophils, eosinophils, basophils, lymphocytes, monocytes), with an accompanying scatter plot of these values across donors.

Supplementary Table 3.

Top marker genes for each of the ten epithelial and stromal cell populations in the combined healthy and tumor atlas, one sheet per population (AT2, Basal 1, Basal 2, Cancer epithelial, Endothelial, Goblet, Ionocytes, Multiciliated, Myofibroblasts, Secretory), each versus all remaining cells, with adjusted P value, log2 fold change,

Wilcoxon score, percentage dropout within and outside the population, mean log-normalized and raw-count expression.

Supplementary Table 4.

Gene set enrichment analysis results for each major epithelial/stromal cell type in the combined atlas. Preranked GSEA results (GO Biological Process 2023, KEGG 2021, and MSigDB Hallmark 2020 collections) for eight cell types (Basal cells, Suprabasal cells, Goblet cells, Secretory cells, Multiciliated cells, Cancer epithelial cells, Myofibroblasts, Endothelial cells), with enrichment score, normalized enrichment score, nominal P value, FDR q value, FWER P value, tag percentage, gene percentage and leading-edge genes for every gene set tested.

Supplementary Table 5.

Ranked list of 1,274 genes differentially expressed between cancer epithelial cells and healthy basal/suprabasal cells. Rank, gene symbol, direction (399 up in cancer, 875 up in healthy), mean log2 fold change across the six tumors, mean -log10(P value) and a combined ranking score (|log2FC| x -log10P).

Supplementary Table 6.

Gene set enrichment analysis (GSEA Preranked, MSigDB Hallmark collection) comparing LUSC cancer epithelial cells against healthy basal cells. All 50 Hallmark gene sets, each labeled by direction (29 up in cancer, 21 up in healthy), with enrichment score, normalized enrichment score, nominal P value, FDR q value, FWER P value, rank at max, and leading-edge genes.

<u>Supplementary Table 7.</u>

Top marker genes for five epithelial lineages within the healthy-donor-only dataset. One sheet per lineage (Basal 1, Basal 2, Goblet, Multiciliated, Secretory), each versus all remaining healthy cells, with adjusted P value, log2 fold change, Wilcoxon score, percentage dropout and mean log-normalized expression.

<u>Supplementary Table 8.</u>

The full list of transcription factors defining each of the four basal subpopulations used in Figures 3 and 4 (Basal I-IV), one column per subpopulation (Basal I, Basal II, Basal III, Basal IV.

<u>Supplementary Table 9.</u>

Gene set enrichment analysis for each basal subcluster, Basal I through Basal IV. Preranked GSEA results per subcluster, with enrichment score, normalized enrichment score, nominal P value, FDR q value and FWER P value for every Hallmark gene set tested.

Supplementary Table 10.

Gene set enrichment analysis results for bulk RNA sequencing of BCi-NS1 cells after HLF overexpression versus control. Panel A: RB1-target and cell-cycle gene sets (9 sets). Panel B: the complete MSigDB Hallmark collection (50 sets). Each row gives normalized enrichment score, P value, false discovery rate, gene set size and leading-edge genes.

Supplementary Table 11.

Research reagents used in this study.

Supplementary Table 12.

Differential expression of each LUSC tumor versus healthy basal/suprabasal cells, all detected genes. Two-sided Wilcoxon rank-sum test for each of the six tumors (SCC1–SCC6) against pooled healthy basal/suprabasal cells, for all 36,601 detected genes.
