## Supplementary methods for "HLF suppresses DREAM target genes and reinstates quiescence in lung squamous cell carcinoma"

The study was approved by the Swedish Ethical Review Authority (formerly Regional Ethical Review Board at Lund University), in adherence to the Declaration of Helsinki (1) (Registration no. 2014/32). All patients provided written informed consent prior to enrolment. Ethical permit numbers: 2014/32, 2017/180, and 2017/519.

Sampling and primary cell processing

Healthy tissue pieces were taken as biopsies from the airway region between branching generations 4–6 and were sequenced individually. Tissue was collected in DMEM/F12 media and dissociated immediately. Tissue pieces were digested, using the solutions within the Miltenyi human tumor dissociation kit (Miltenyi), in the c-tubes connected to the Miltenyi gentleMACs Octo Dissociator (Miltenyi) with heaters at 37 °C. Antibiotics + antimycotics (Penicillin, streptomycin, and gentamicin) were added throughout. After dissociation, the suspension was passed through a 70µm cell strainer. The cell suspension was then centrifuged and red blood cells were lysed from the cell pellet using red blood cell lysis buffer. The remaining cells were frozen in liquid nitrogen for future storage and subsequent sequencing. Cancer tissue was received as explanted tissue that was stored on ice in cancer cell medium. The tissue was then processed immediately and fresh frozen few hrs after receiving the explanted tissue.

Preparation for single cell RNA seq

Removal of dead cells, doublets and other debris was performed with a magnetic microbead based column dead cell removal kit (Miltenyi). The effluent collected was the viable cell fraction. This was then followed by white blood cell exclusion through CD45 positive magnetic cell separation (Miltenyi). The CD45 negative cell flow through was used for subsequent RNA sequencing. The cells were encapsulated into emulsion droplets as single cells using the Chromium Controller (10X Genomics). scRNAseq libraries were constructed using the Chromium Single Cell Gene Expression 3' v3 Reagent Kit according to the manufacturer's instructions. Reverse transcription and library preparation was performed on a C1000 Touch Thermal Cycler (Bio-Rad). Amplified cDNA and final libraries concentrations were measured with a Qubit 4 Fluorometer (Invitrogen) using the dsDNA High Sensitivity Assay (Invitrogen). cDNA and library traces were evaluated on a TapeStation (Agilent) using High Sensitivity D5000 and D1000 Screen Tapes (Agilent). Individual libraries were diluted to 1.5 nM and pooled for sequencing.

Pre-processing of scRNA-seq data, quality control, normalization, and annotation of single cell RNA-Seq Data

Mapping of the FASTQ files and gene counting were performed using Cell Ranger pipeline (v6.1, 10x Genomics) with GRCh38 human reference genome. Secondary pre-processing and downstream analysis were done within the Scanpy ecosystem. Cells with fewer than 2,000 detected UMIs or with >12% of UMIs derived from mitochondrial genes were excluded (Supplementary Figure E1); exceptions to quality control filtering were allowed for ionocyte cell subpopulation. Putative doublets were identified and excluded using Scrublet algorithm (2) Gene expression values were log-transformed and normalized to the median expression across the dataset. The top 3,000 highly variable genes (HVGs) were selected to compute 50 principal components, which were used to construct a batch-balanced k-nearest neighbors (BBKNN) graph (3). Uniform Manifold Approximation and Projection (UMAP) was applied for two-dimensional visualization; Leiden clustering algorithm was used to compute cell clusters. Cell type annotation was performed at the cluster level based on known marker genes for lung epithelial cell types. Additionally, CellTypist models Human Lung Atlas (4) and Human IPF (5) were used to provide a second opinion for each cell and for refinement of the cluster-based annotation. The data was additionally mapped to the HCLA to confirm our findings recapitulating established markers of lung epithelial subtypes (Supplementary Figure E1, E3 and E4).

Differential expression analysis was done using a two-sided Wilcoxon rank-sum test for the following comparisons: cell type-specific (each cell type versus all others, within healthy samples); age-related (aged versus young, within each cell type in healthy samples); condition-specific (cancer epithelial cells versus healthy epithelial cells), and patient-specific (cancer epithelial cells from each patient versus healthy epithelial cells). Gene Set Enrichment Analysis (GSEA) using MSigDB gene sets was conducted using 1000 gene set permutations for genes ranked by Wilcoxon score. As complementary validation, over-representation analysis (ORA) for genes with adjusted p-value $< 0.05$ and absolute log fold change $> 0.15$ was performed using the same gene set collections. For pseudotime analysis, data from healthy samples were re-integrated separately. Diffusion pseudotime was computed using a root cell (Supplementary Figure E10) defined by the highest expression of TP63 within the basal cell cluster. Additionally, a PAGA-graph on high-resolution clusters was computed to verify pseudotime at cluster level and to evaluate potential branching. Gene regulatory network (GRN) was inferred using SCENIC tool (6). Known transcription factors (TFs) and the top 3,000 HVGs were used as input. Gene co-expression matrix was calculated with GRNBoost2 algorithm and regulon activity was inferred using transcription factor motif enrichment within 10 kb up- and downstream regions of transcription start sites.

Clustering of the basal compartment into four pseudotime-resolved states

Conserved differentially expressed genes (DEGs) between cancer epithelial cells and healthy Basal 1 and Basal 2 cells were defined as genes showing significant differential expression with a consistent direction of log fold change in 5 of the 6 six tumor samples. Within the healthy airway trajectory, basal cells were resolved into four transcriptionally distinct states (Basal I–IV, corresponding to the quiescent, activated, hyperactivated and primed populations described in the main text) using Leiden clustering at resolution 1.5. Marker genes for each basal state were identified using a two-sided Wilcoxon rank-sum test comparing each basal cluster against the remaining three (Supplementary Table 7) and were correlated against one another to determine similarity and possible transition (Supplementary Figure E11). To confirm this four-state structure, the combined healthy and cancer dataset was subset to the basal compartment and re-clustered; the resulting clusters were matched to the original Basal I–IV labels by nearest-neighbor label transfer in a shared principal component space.

Disruption of the DREAM-dependent transcriptional program was assessed using 210 curated p53–p21–DREAM–CDE/CHR target genes defined by Fischer and Engeland [8]. For each gene, the mean log2 fold change across the six tumors versus healthy Basal cells was calculated; genes with a mean log2 fold change $\geq 0.5$ were classified as upregulated, $\leq -0.5$ as downregulated, and all others as unchanged (Supplementary Figure E6 and E7). Gene symbols revised since the original publication were updated to current HGNC nomenclature prior to classification (e.g., C11orf82→DDIAS, CASC5→KNL1, GSG2→HASPIN). The same approach was applied to curated gene sets for hypoxia/Warburg metabolism, immune/MHC signaling and the NRF2 antioxidant response, in addition to other gene sets of interest such as the mitotic spindle etc (Supplementary Figure E8).

Data and code availability

All data has been uploaded to the National Center for Biotechnology Information (Gene Expression Omnibus accession number: GSE295552. The analysis pipeline and the code will be available at: github.com/SofiaFreiman/Lung_epitelium/

### Cell culture

All cells were grown as submerged cultures for this study. The immorlatlized BSC cell lines; BCi-NS1 (6) was kindly provided by the Crystal lab, while primary BSC (HUBEC) were retrieved from bronchoscopy, and the two $KRT5^{+}$ lung squamous cell carcinoma cell lines; HCC-95 and HCC-1588 (Cancer Research Institute, Seoul National University College of Medicine, Korean Cell Line Bank). The cell lines were grown in 2D according standard protocols for growing basal epithelial cells using the PneumaCult growth media supplemented with 0.5% antibiotics/antimycotics, 0.5% hydrocortisone and the PneumaCult growth medium. The cancer cells were grown in DMEM medium supplemented with 10% FBS and 0.5% antibiotics/antimycotics. All cells were plated on collagen-coated tissue culture plates and allowed to reach 70% confluence before they were trypsinized. At 70% confluence, cells were passaged by dislodging with trypsin or TrypLE and replated at a 1:4 split. Media was replaced every 2-3 days depending on the confluence of the cells.

### Virus Production

Human embryonic kidney (HEK) 293T cells were used to package the produced lentiviral particles. Before transfection, 7 million cells were seeded in a 15-cm dish to achieve approximately 80% confluence after 1 day. The following day, cells were transfected as by using 10 µg of lentiviral vector, 7.5 µg of psPAX2.G lentiviral packaging vector, and 2.5 µg of pMD2 envelope vector were combined with 60 µl of polyethylenimine (1 mg/ml) in 2 ml of Opti-MEM and incubated for 15 min at room temperature. The PEI-DMEM mixture was added dropwise to HEK293T cells and, 12 hours later, replaced with fresh medium. Viral supernatants were collected 48, 60, and 72 hours after transfection, filtered through a 0.45-µm cellulose acetate filter, and concentrated with the Lenti-X Concentrator. Overexpression was induced via transduction of cells with lentiviral particles in the presence of polybrene (8 µg/ml).

Flow cytometry

For proliferation analysis, GFP positive cells were analyzed, DAPI was used to exclude dead cells. All experiments were performed using FACS Canto, FACS A1, and FACS LSR II cytometers (Becton Dickinson) and analyzed by Flow Jo software (Tree Star, version 9.5.1 or 10.0.2)

Flow cytometric cell cycle analysis

Harvested cells were resuspended in 1mL containing Hoeschst (1mg/mL) and incubated at $37^0$C for 1hr. this was followed by the addition of 5uL Pyronin Y (100 µg/mL) and incubated at $37^0$C for a further 15 mins. Cells were then analyzed using flow cytometry. G0 and G1 cell cycle fractions were distinguished by co-staining transduced and untransduced cells with Hoechst 33342, and Pyronin Y, marking DNA and RNA content respectively. Cells with 2N DNA content and low Pyronin Y signal were gated as G0; cells with 2N DNA content and higher Pyronin Y signal were gated as G1.

Quantitative real-time PCR (qRT-PCR)

E2F1 and the DREAM-CDE/CHR target genes CDC20 and CCNB2 were measured by qRT-PCR (QuantStudio Rt-PCR system (Thermo-Fisher)) 24 hours after HLF or control lentiviral transduction in HCC95 and HCC1588 cells, across three paired experiments per cell line (Fig. 6N). RNA was extracted according to the RNeasy Micro Kit Qiagen after lysis with RLT buffer supplemented with β-marcaptoethanol. The extracted RNA was prepared for cDNA synthesis using the Invitrogen Superscript IV on the C1000 Touch thermal cycler (Bio-Rad). Housekeeping genes UBC and RPLP0 were used to normalize the data. The results were quantified using the ΔΔCt method.

Bulk RNA sequencing of HLF-overexpressing cells

BCi-NS1 cells transduced with HLF-IRES-ZsGreen lentivirus or control vector were profiled by bulk RNA sequencing 48 hours after transduction to identify genes suppressed by HLF overexpression (Figure 5H, I, J). Differential expression results (log2 fold change, HLF-overexpressing versus control) were used directly as a processed table, consistent with standard DESeq2-style output. Gene-level counts were obtained by transcript quantification with Salmon (7), summarised to gene level with tximport (8) using length-scaled TPM. Estimated counts were rounded to integers and analysed in R (v4.3.1) with DESeq2 (v1.42.0) (9). Size factors were estimated with the median-of-ratios method and dispersions were modelled with a parametric fit. Differential expression was assessed using two-sided Wald tests. Log2 fold changes are reported without shrinkage. P-values were adjusted with the Benjamini–Hochberg method, and genes with an adjusted p-value $< 0.1$ were considered differentially expressed. For visualisation, counts were transformed with DESeq2's variance-stabilising transformation (VST). Results were displayed as row-scaled (z-score) heatmaps using pheatmap.

Trypan blue cell proliferation assay

Transduced lung Squamous carcinoma cell line HCC-95 and HCC-1588 were then sorted into pure populations using FACS. The cells were then placed in 2D submerged culture and allowed to grow until the cells reached 70% confluence. The cells were then passaged as above. During the passage and re-suspension, cells were stained with 0.4% trypan blue dye and live cells were counted on a hemocytometer. The cells were categorized into NV (no vector), CV (control vector), and HLF (over-expression vector) groups and were seeded in a 12 well plate at a cell density of 5000 cells per well. Cells were maintained under standard culture conditions with incubation periods of 4 and 8 days, to assess growth kinetics.

Air-liquid interface culture and differentiation

To assess the viability of using the BCi-NS1.1 cell line we wanted to test whether the cell line would be able to recapitulate the differentiated airway epithelium. Approximately 80,000 cells were plated per insert were seeded onto 0.4 μm pore-size Polyethylene terephthalate (PET) transparent membrane inserts (24-well format) and maintained in PneumaCult basal media under submerged conditions for 4 days at 37°C with 5% $CO_2$. Post-confluence, an air-liquid interface (ALI) was established by aspirating the apical medium and replacing the basolateral medium with PneumaCult-ALI differentiation media, supplemented with retinoic acid and growth factor cocktails. Cells were cultured under ALI conditions for 28 days, with differentiation media replenished every 48 hours. Weekly mucus clearance was performed by gentle aspiration of the apical surface using sterile PBS. For the histological processing after 28 days of ALI culture inserts were made into paraffin chips according to the protocol from (10) Membranes were carefully excised, trimmed, and subjected to serial dehydration in graded followed by clearing in xylene (15 minutes) and paraffin embedding using a standard tissue processor. Embedded blocks were sectioned at 10 µm thickness using a rotary microtome (HM 325, Thermo Scientific). The sections were then processed for immunostaining (Supplementary Fig. E16).

Spheroid formation, longitudinal imaging and morphometric analysis

HCC1588 and HCC95 squamous cell carcinoma cells expressing an HLF-overexpression vector (OV), control vector (CV), or no vector (NV) were cultured as spheroids using a suspension culture protocol adapted from (11). Cells were harvested by trypsinization, counted, and seeded at 1,000 cells/well in 200 µL medium in 96-well low-adherence plates, with ten replicate wells initially established per condition. Spheroids were maintained at 37°C and 5% $CO_2$ in PneumaCult-ALI-based medium supplemented with hydrocortisone, heparin, IL-13 (5 ng/mL) and Y-27632 (10 µM). Spheroids were imaged using an Olympus IX70 inverted microscope. Bright-field images were acquired for spheroid morphology and diameter measurements. For fluorescence imaging of live-cell viability staining, Hoechst 33342 was imaged using the DAPI channel, fluorescein diacetate (FDA) using the FITC channel, and propidium iodide (PI) using the Cy5 channel. Hoechst was used to visualize nuclei, FDA to identify metabolically active/viable cells, and PI to identify cells with compromised plasma membranes. For quantitative morphometric analysis, five independently seeded spheroids per condition (n = 5 wells) were analyzed from bright-field images using FIJI/ImageJ. Four diameters passing through the central spheroid cell mass, horizontal, vertical, left diagonal and right diagonal were measured for each spheroid and averaged to obtain its mean diameter. Comparisons were performed between vector conditions within each cell line. Quantitative size analysis was restricted to the early culture period because prolonged culture resulted in increasingly irregular spheroid morphology and development of an apparent non-viable central cell mass (Figure 6L, M).

Patient-derived LUSC organoid culture, lentiviral transduction, and spheroid growth assays

Tumor tissue from a patient with suspected lung cancer, subsequently diagnosed as squamous cell lung cancer, was collected for research purposes during routine diagnostic bronchoscopy. The sample was immediately put in Advanced DMEM/F-12 medium supplemented with 1M HEPES Buffer, 1X GlutaMAX, and 1X Antibiotics / Antimycotics and transported to the research facility within 3 hours from the time of sampling. Upon arrival at the research facility, the sample was processed, cultured, and maintained as described (12), but the TrypLE digestion step was skipped, and reagent volumes were scaled down to account for the small size of our sample.

After tissue collection, the sample was enzymatically dissociated and incubated in 0.5 ml pre-warmed digestion medium at 37 °C with gentle agitation for 30 min to 1.5 h. Every 15 min the tissue was vortexed briefly and checked under the microscope. The digest was passed through a 100 µm cell strainer. Cells were pelleted at 300–600 × g for 5 min at 4 °C. Pellets were resuspended in 20 µl organoid culture medium and mixed on ice with 40 µl Matrigel, giving a final ratio of about 30–35% medium to 65–70% Matrigel. A dome of 50–60 µl were dispensed into a pre-warmed 24-well plate. After 1 min the plates were inverted and incubated at 37 °C for 30–40 min until the domes had set. Each well was then gently overlaid with 600 µl pre-warmed complete organoid medium containing Y-27632.

The primary LUSC tumor organoids were then transduced with HLF-IRES-ZsGreen or control lentivirus. Transduced organoids were assessed for perimeter and size-class distribution (Fig. 6O, P) and retained TP63 expression by immunofluorescence (Fig. 6Q; Supplementary Fig. E22).

Histopathology

Where needed, samples were fixed overnight at 4 °C in 10% neutral buffered formalin (organoid staining). Following fixation, samples underwent graded ethanol and isopropanol dehydration series (Fisher Scientific) before paraffin embedding. 5µm thick sections were cut with a microtome (Epredi HM 355S Automatic Microtome) and placed on SuperFrost Plus glass slides and allowed to dry. Prior to staining the slides were placed in a 65 °C oven for deparaffinization. Chemical deparaffinization was performed through incubations of the tissue sections in 2x Xylene 100%, followed by a consecutive descending series of Ethanol 2 x 100%, 95%, 70% and 50% and finally a rinse in PBS. Each step was performed for 2 minutes. Then slices were placed in citrate buffer and heat-induced antigen retrieval was performed using the PTlink Dako (Agilent). Then tissue sections were permeabilized with 0.1% Triton X-100 and 1% BSA in PBS for 15 min. The tissue sections were blocked in 5% BSA in PBS for 1 hour with gentle agitation at room temperature. Immunolabeling was performed overnight at 4°C with antibodies in 1% BSA serum in PBS, followed by secondary antibody staining for 1 hour at room temperature in 1% BSA serum in PBS. Nuclei were stained with DAPI (5 ug/ml) for 5 minutes at room temperature. The lung tissue sections were rinsed in PBS and mounted with fluorescence mounting medium.

TCGA expression and DNA methylation analysis

HLF transcript abundance, HLF promoter methylation, and overall survival stratified by HLF expression were analyzed in the TCGA lung squamous cell carcinoma (LUSC) cohort (n = 503 primary tumor, n = 52 normal methylation: n = 370 primary tumor, n = 42 normal; survival: n = 247 low-HLF, n = 248 high-HLF).

Statistics

Data were subjected to a normality test before using one-way analysis of variance (ANOVA), two-way ANOVA, Kruskal-Wallis, or Mann-Whitney test. Group comparisons were corrected using Šidák's, Dunn's, or Tukey's multiple-comparison test with GraphPad Prism version 9 (GraphPad). Alternatively, a Wilcoxon rank-sum test was performed using the "wilcox.test" function in R software. P values are shown when relevant (*P < 0.05, **P < 0.01, ***P < 0.001, and ****P < 0.0001, ns—not significant).
