## Supplementary figures for "HLF suppresses DREAM target genes and reinstates quiescence in lung squamous cell carcinoma"

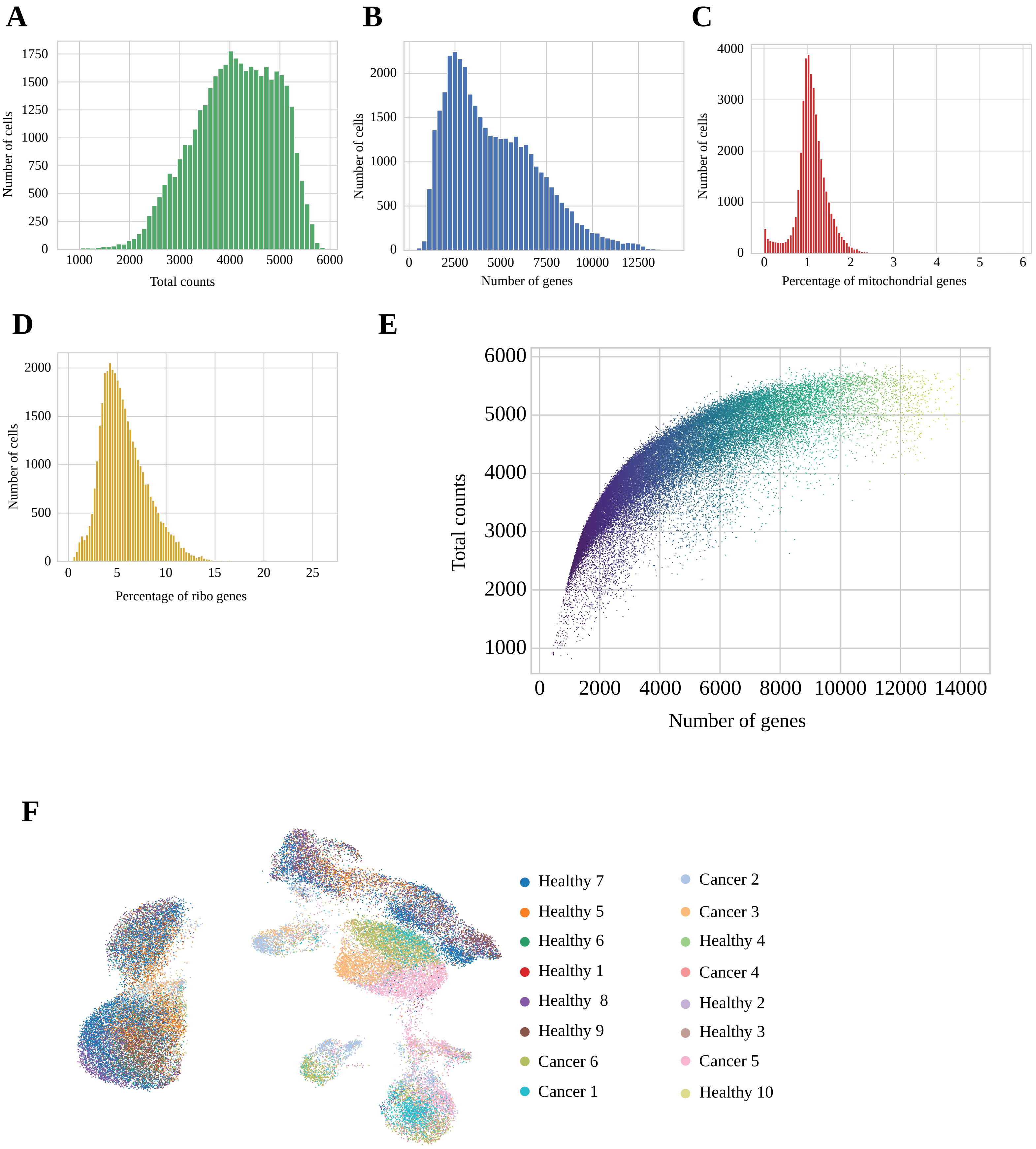


**Supplementary Figure E1**

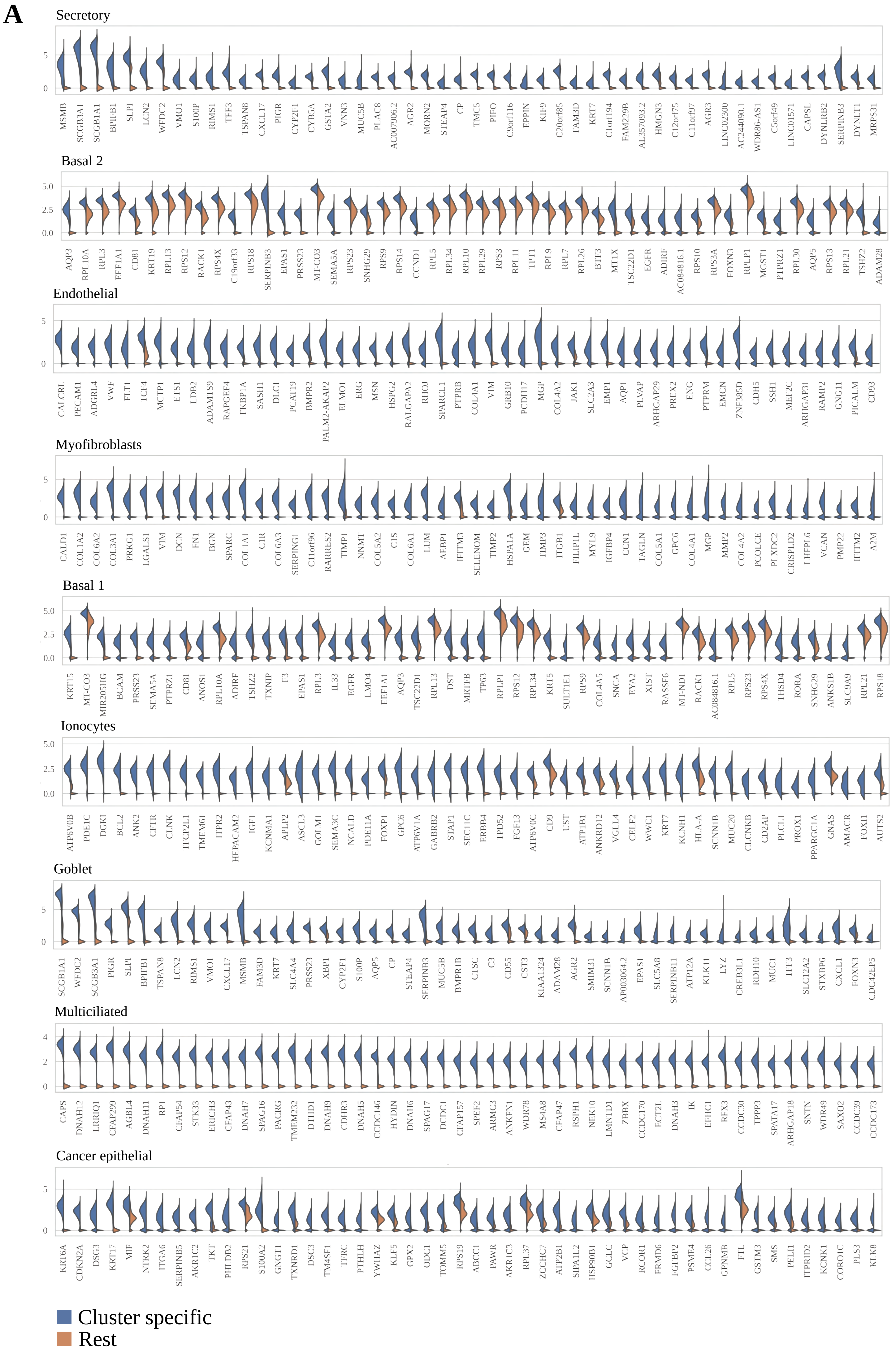


**Supplementary Figure E2**

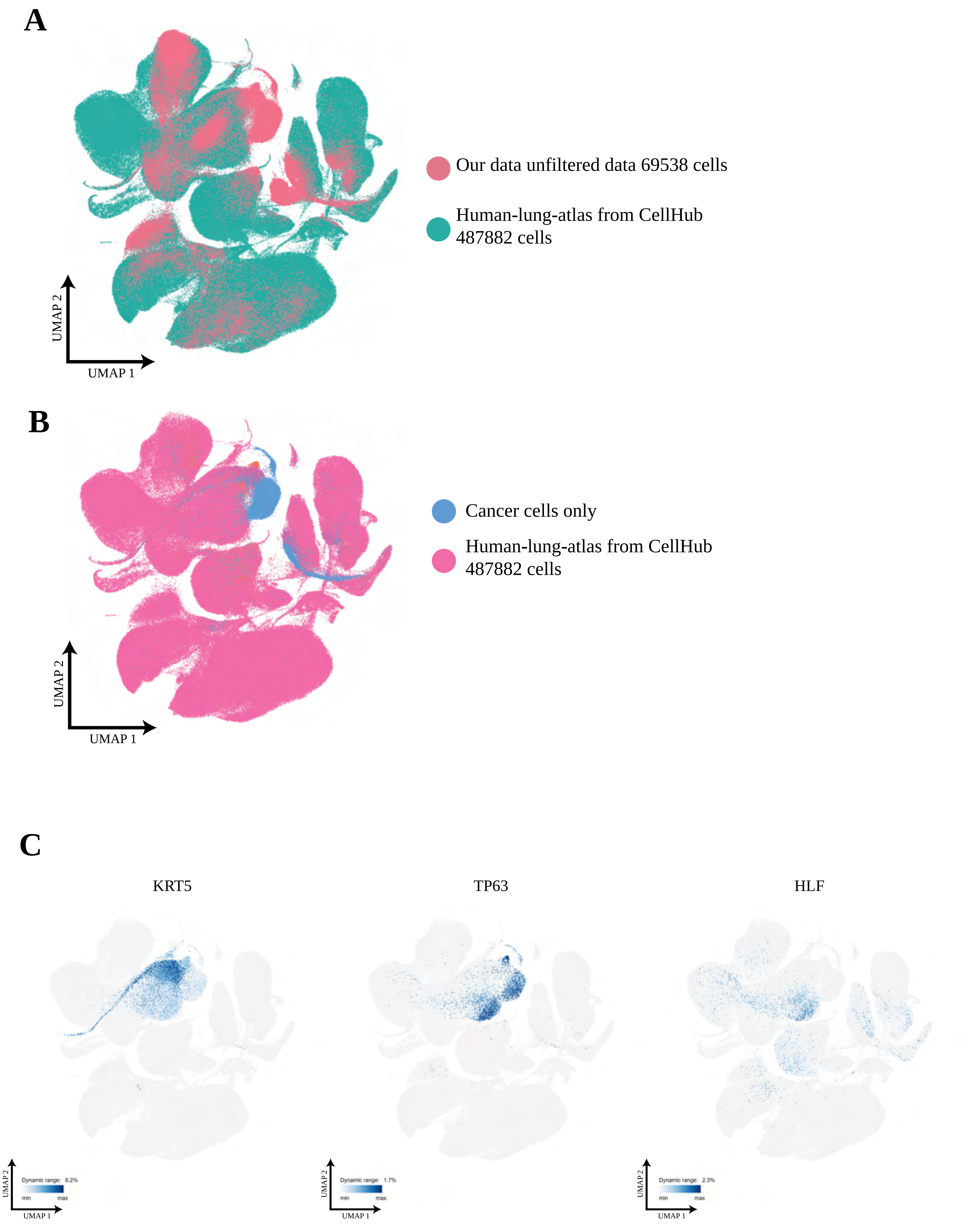


**Supplementary Figure E3**

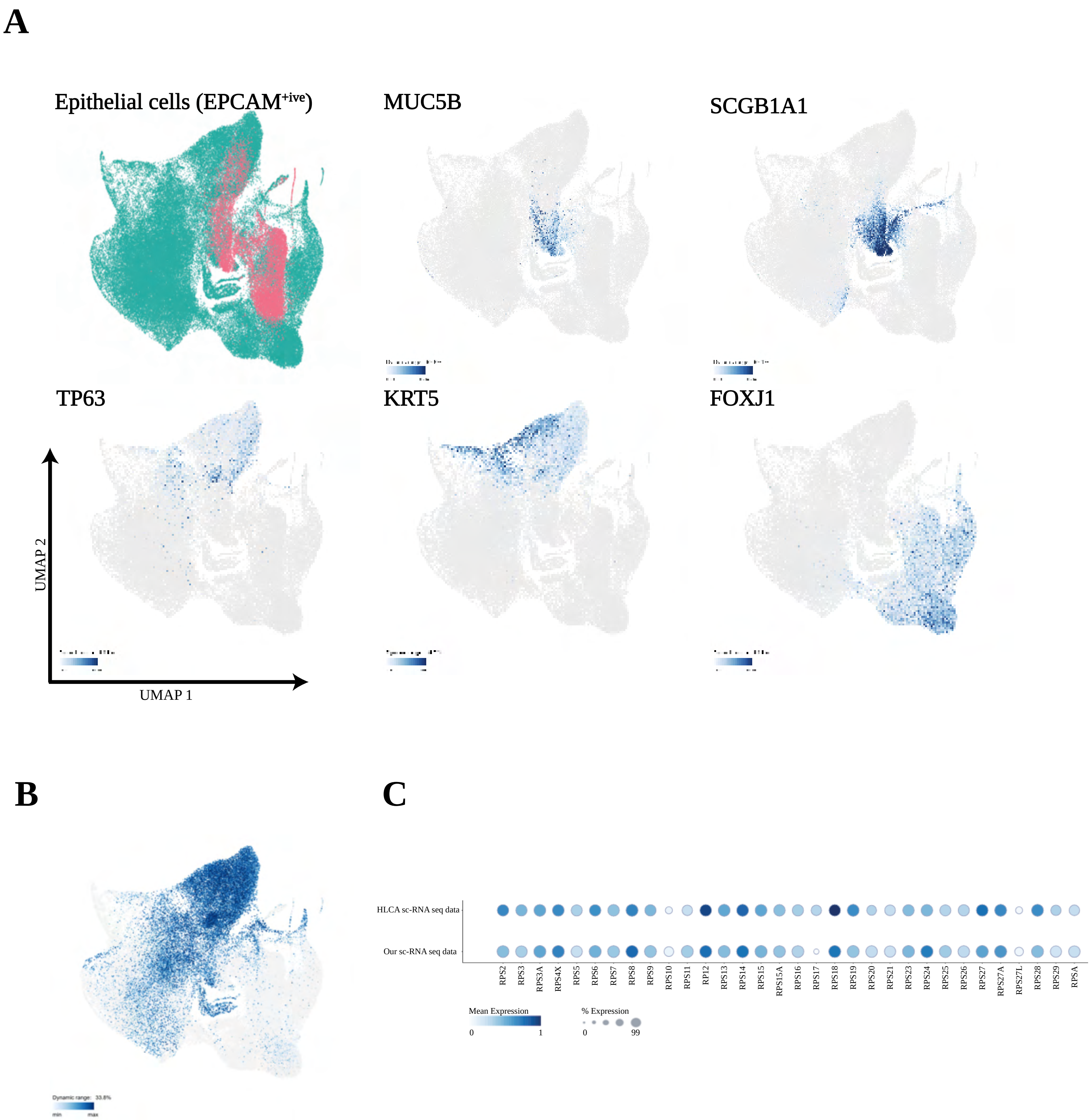


**Supplementary Figure E4**

### Expanded Ribosomal gene expression

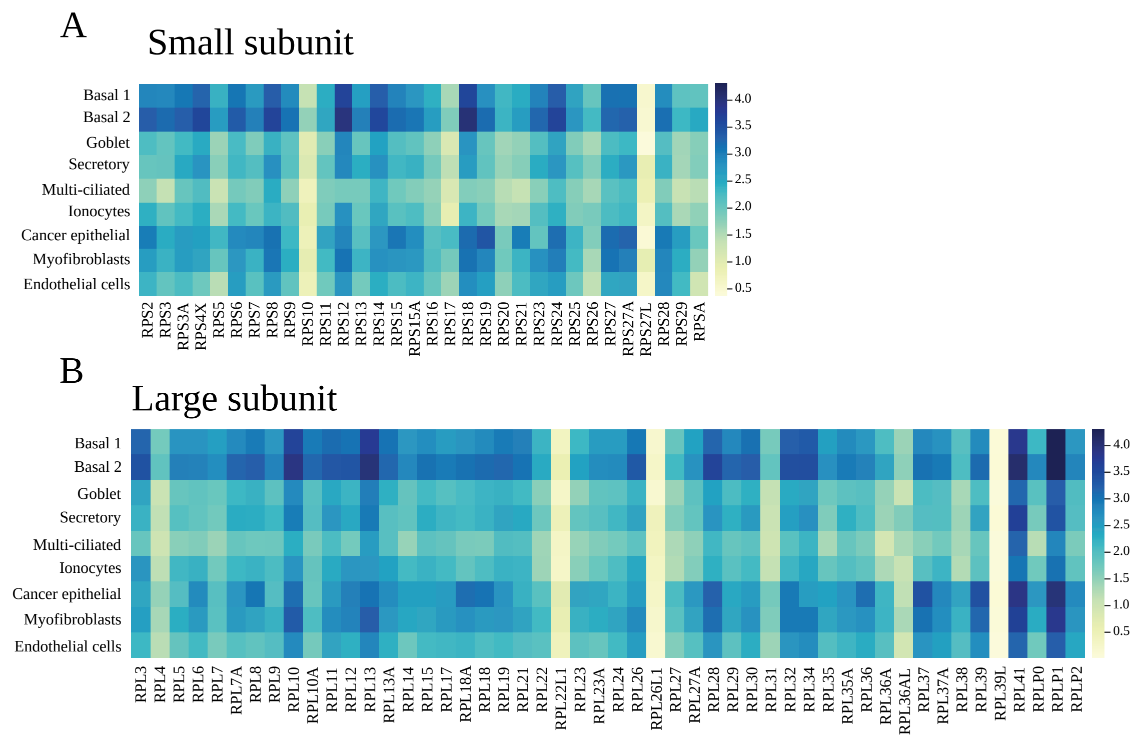


**Supplementary Figure E5**

p53-p21-DREAM Target genes (n=210)
Fischer and Engeland, Nucleic Acids Res, 2016
146 Up (70%) 48 Unchanged (23%) 14 Down (7%)

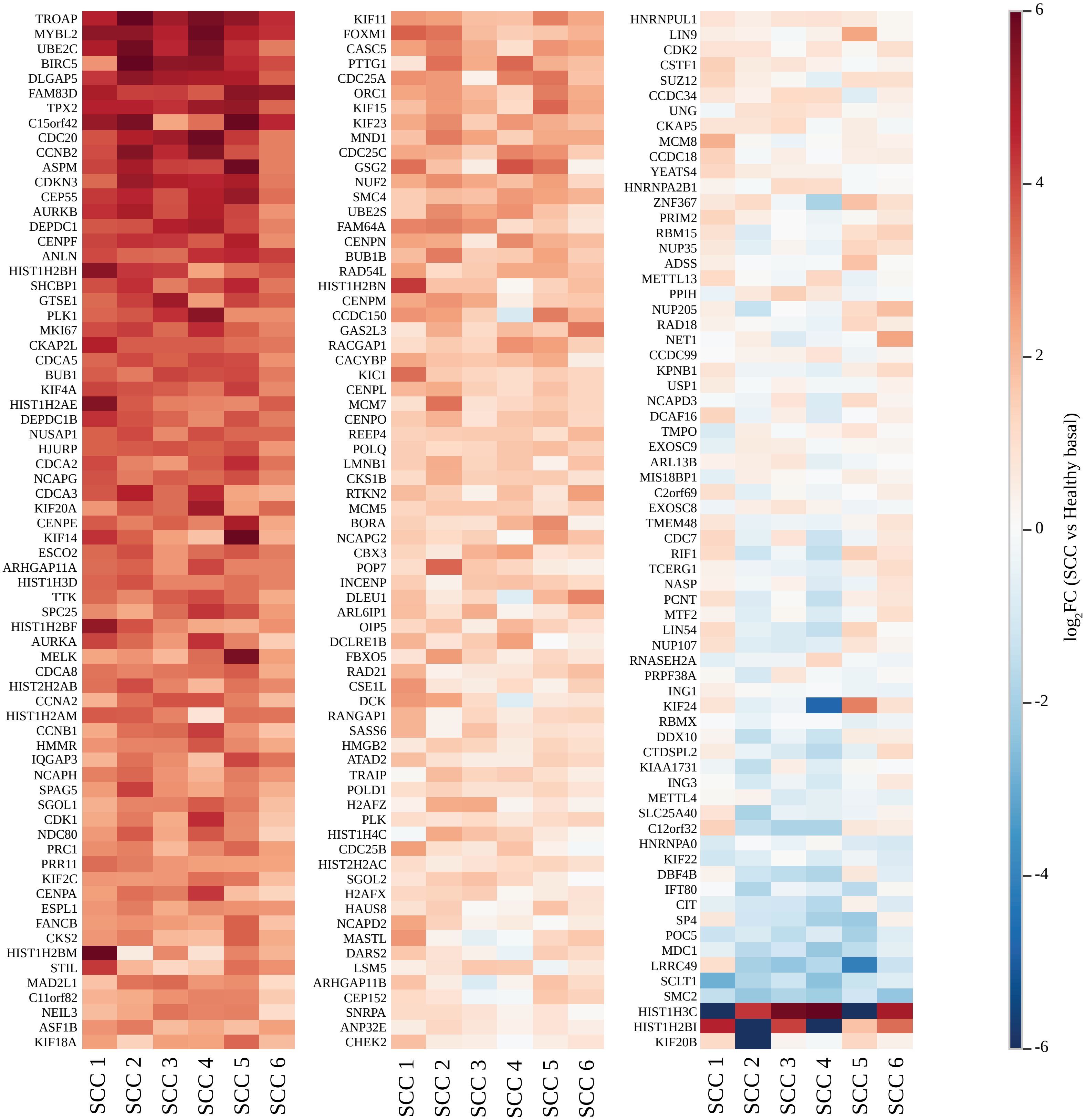


**Supplementary Figure E6**

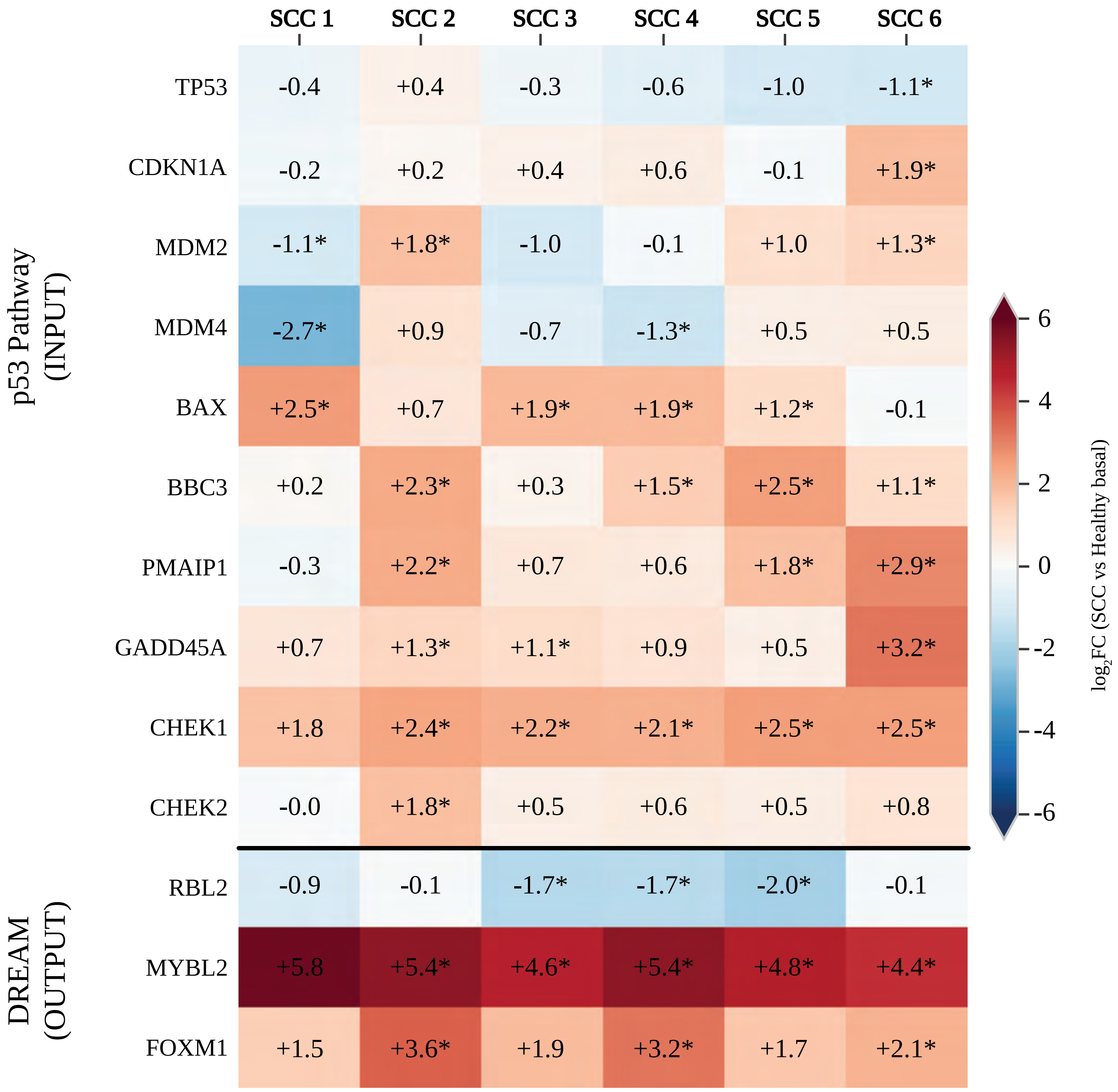


**Supplementary Figure E7**

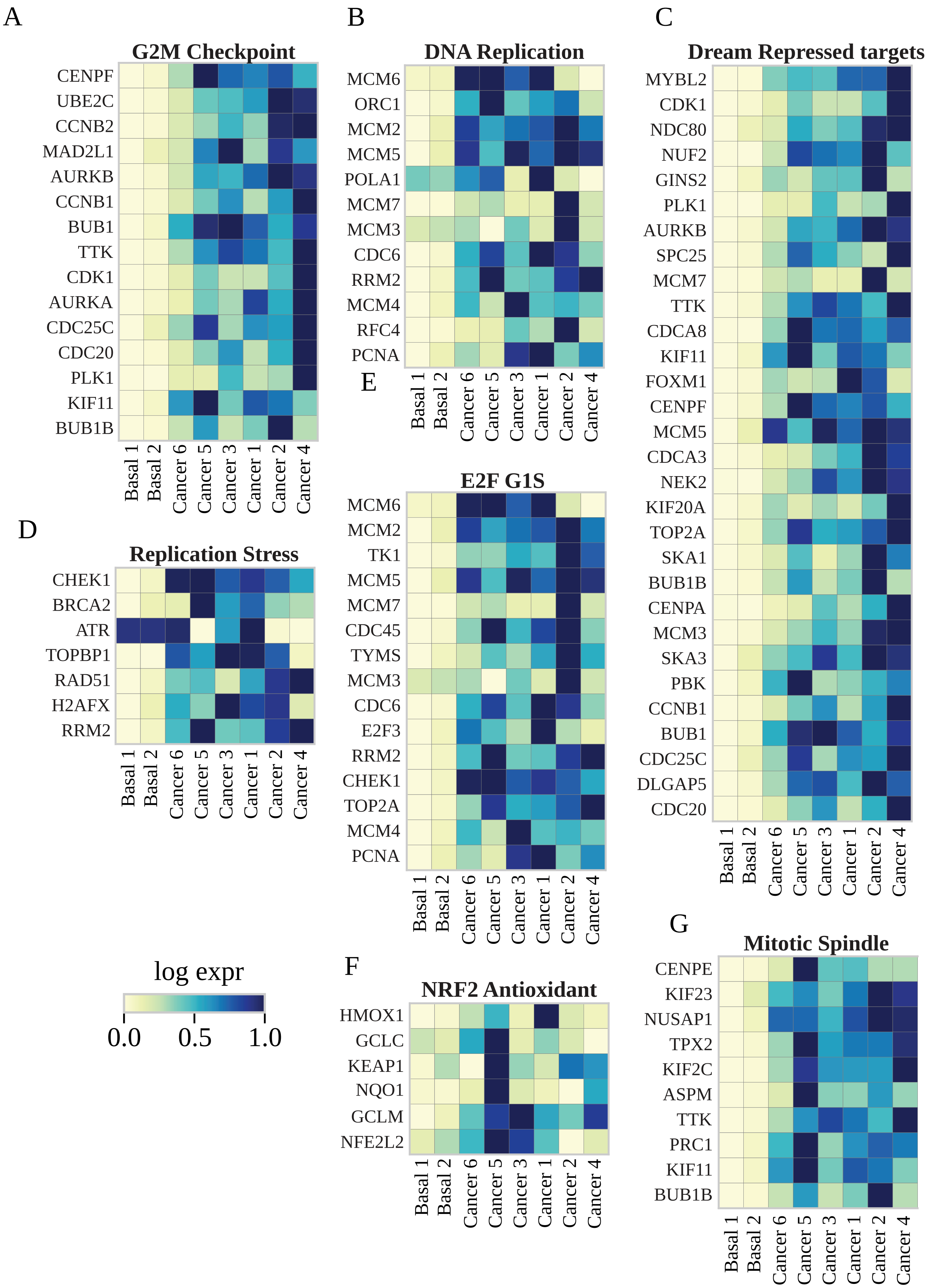


**Supplementary Figure E8**

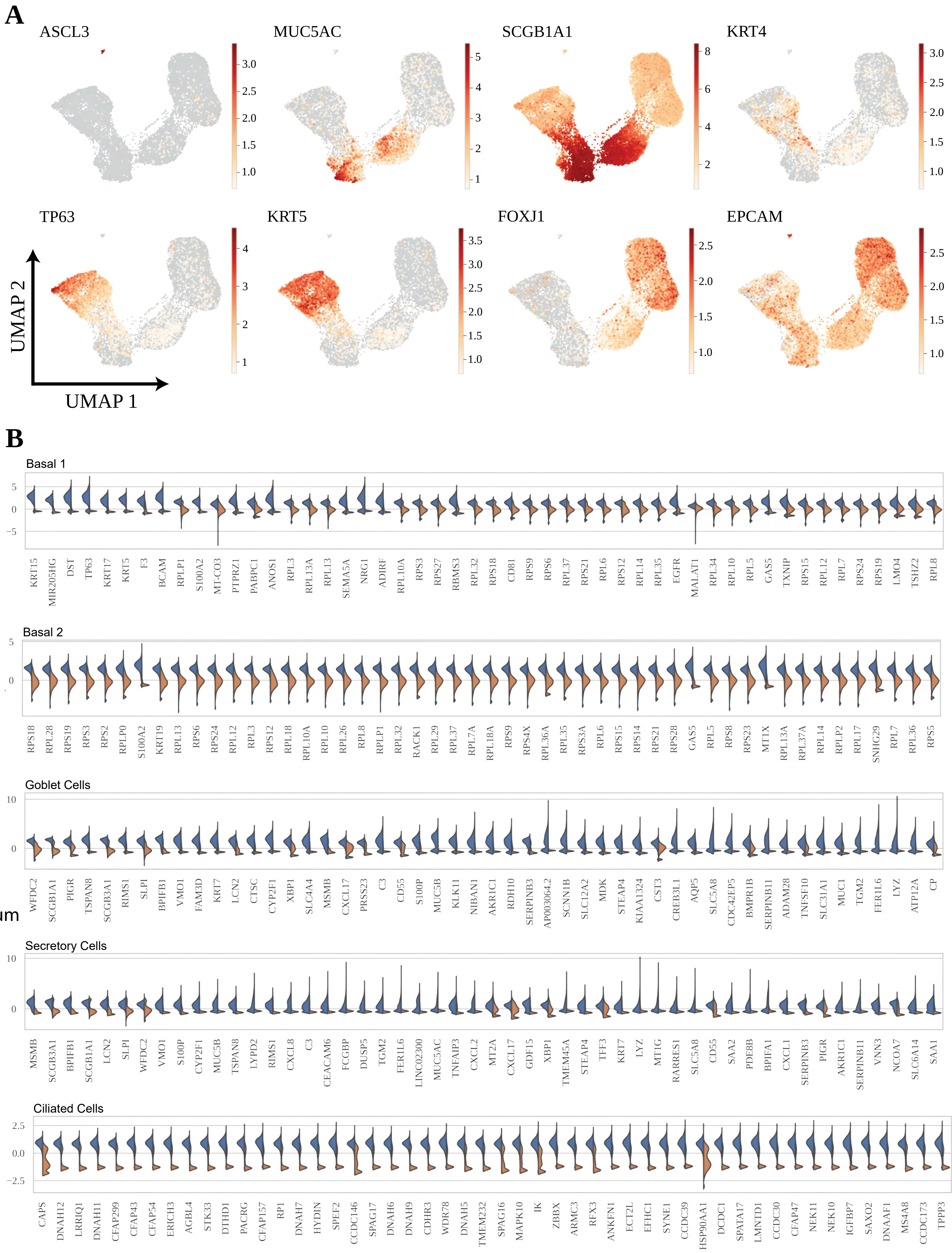


**Supplementary Figure E9**

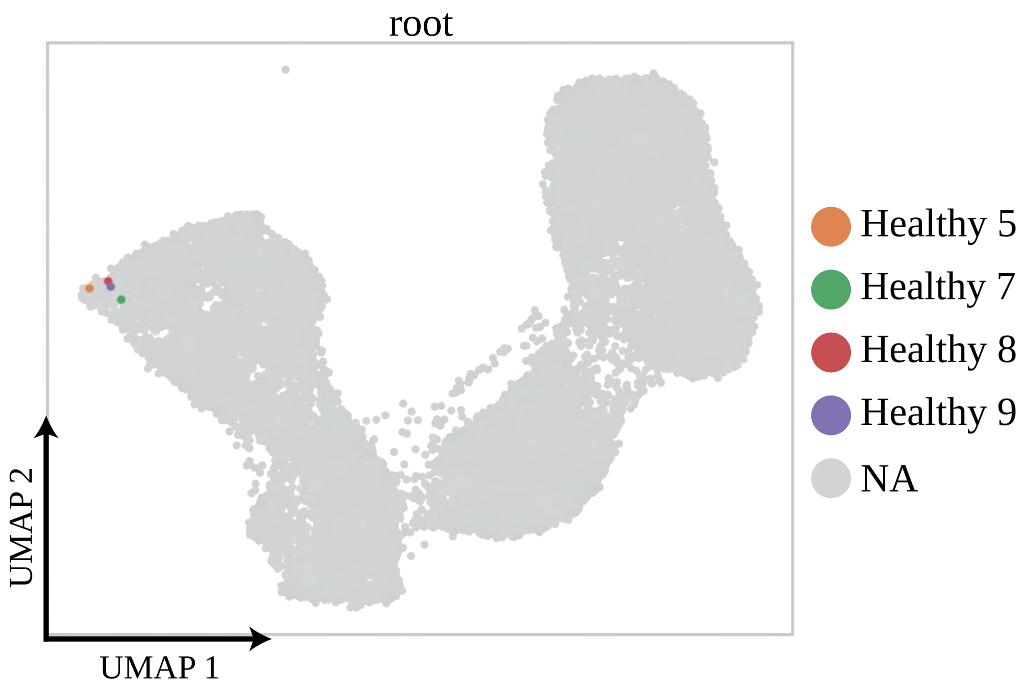


**Supplementary Figure E10**

A

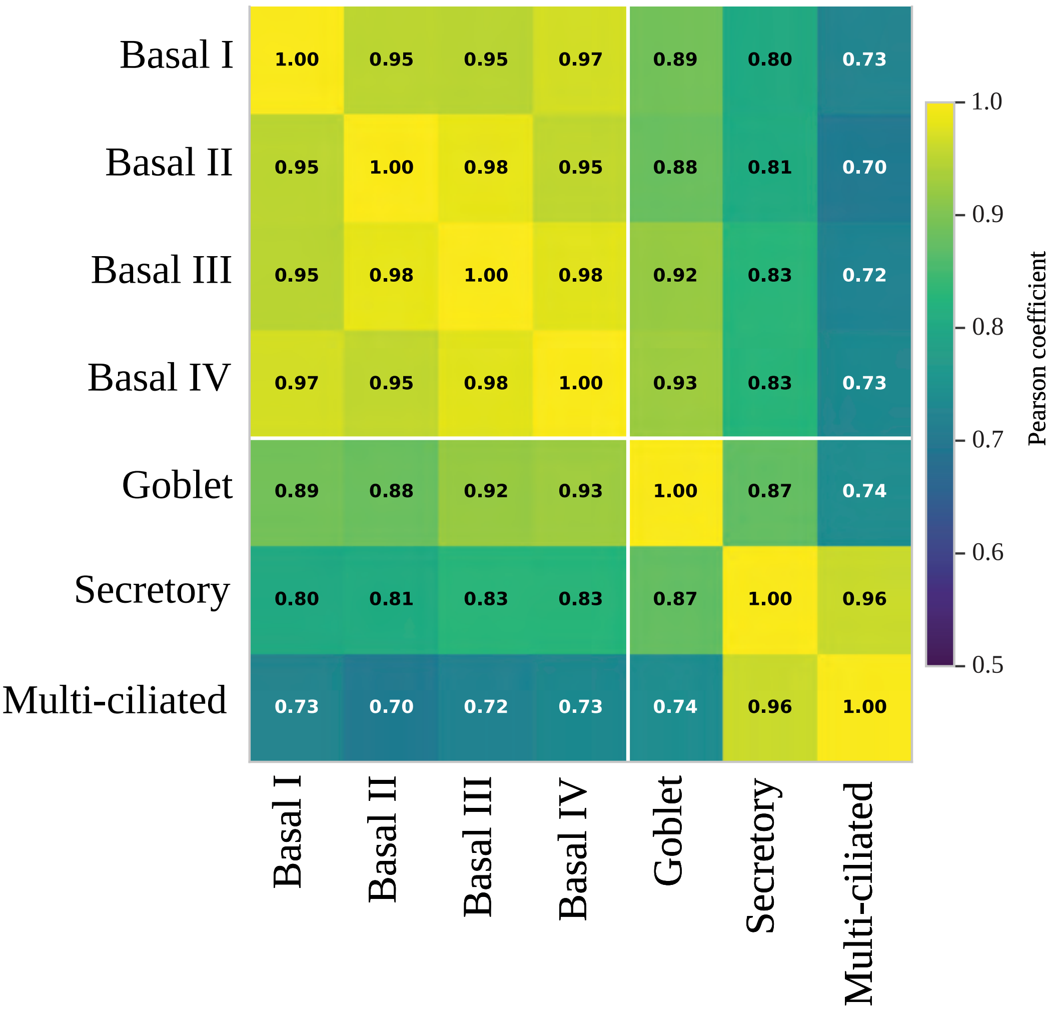


Supplementary Figure E11

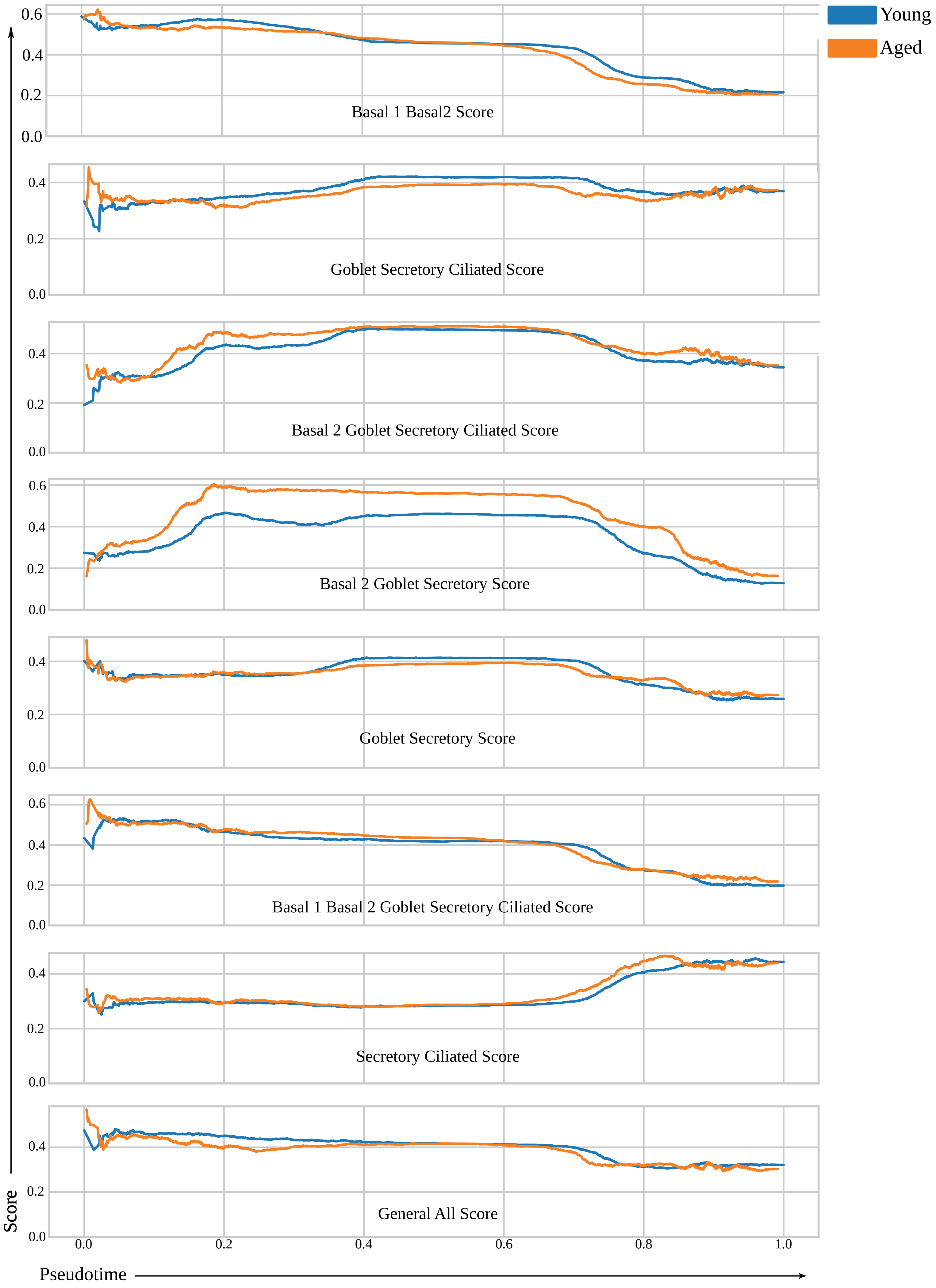


**Supplementary Figure E12**

A

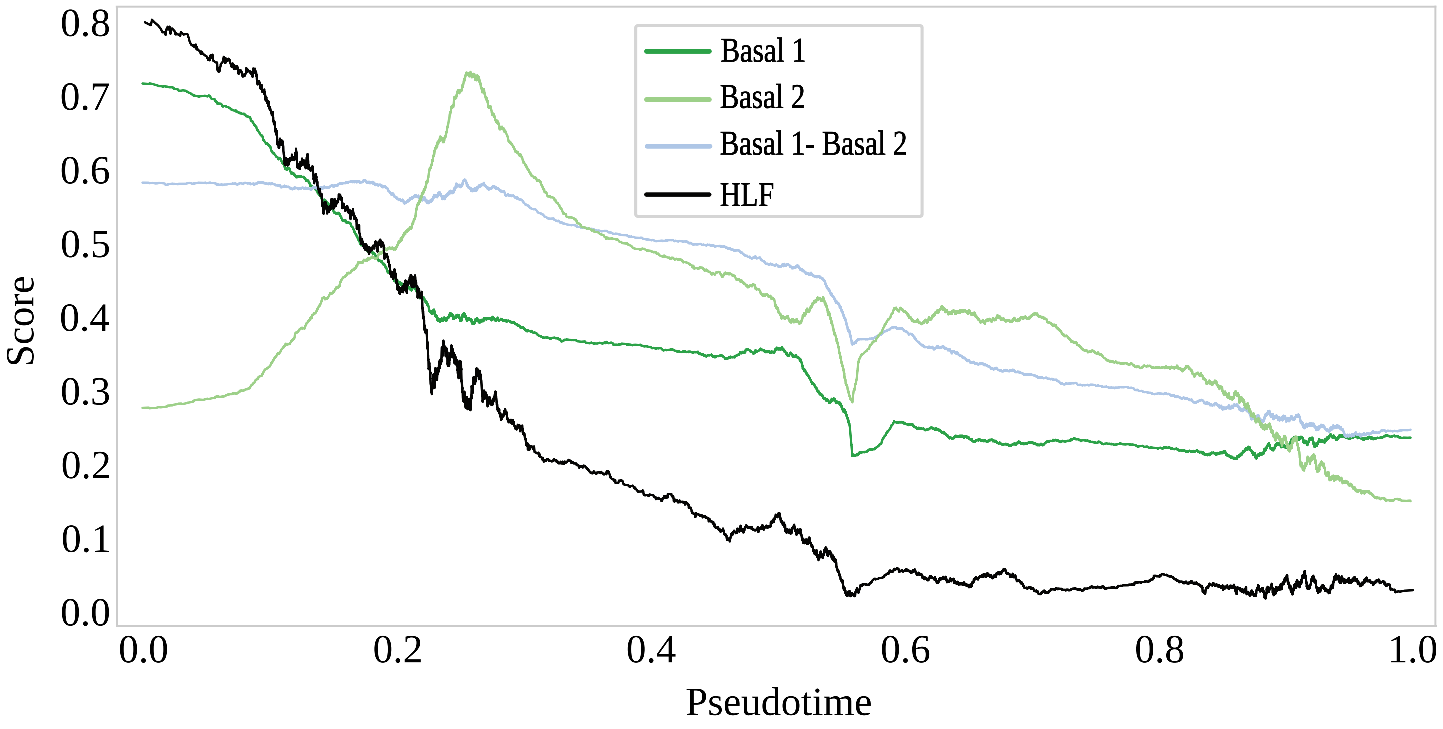


**Supplementary Figure E13**

**Full list of basal relevant transcription factors normalized to each row**

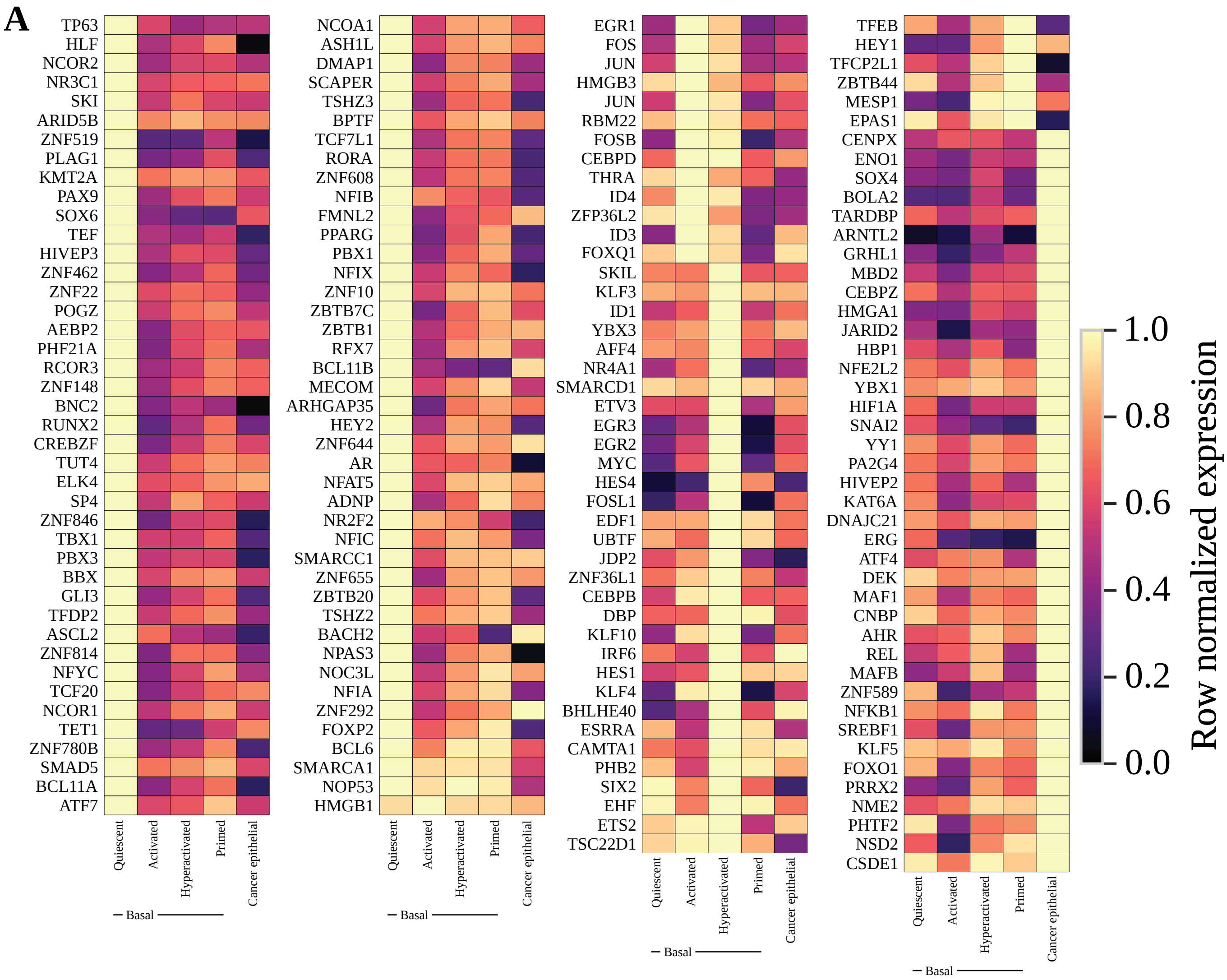


**Supplementary Figure E14**

### Full TF list (% of each gene's own maximum)

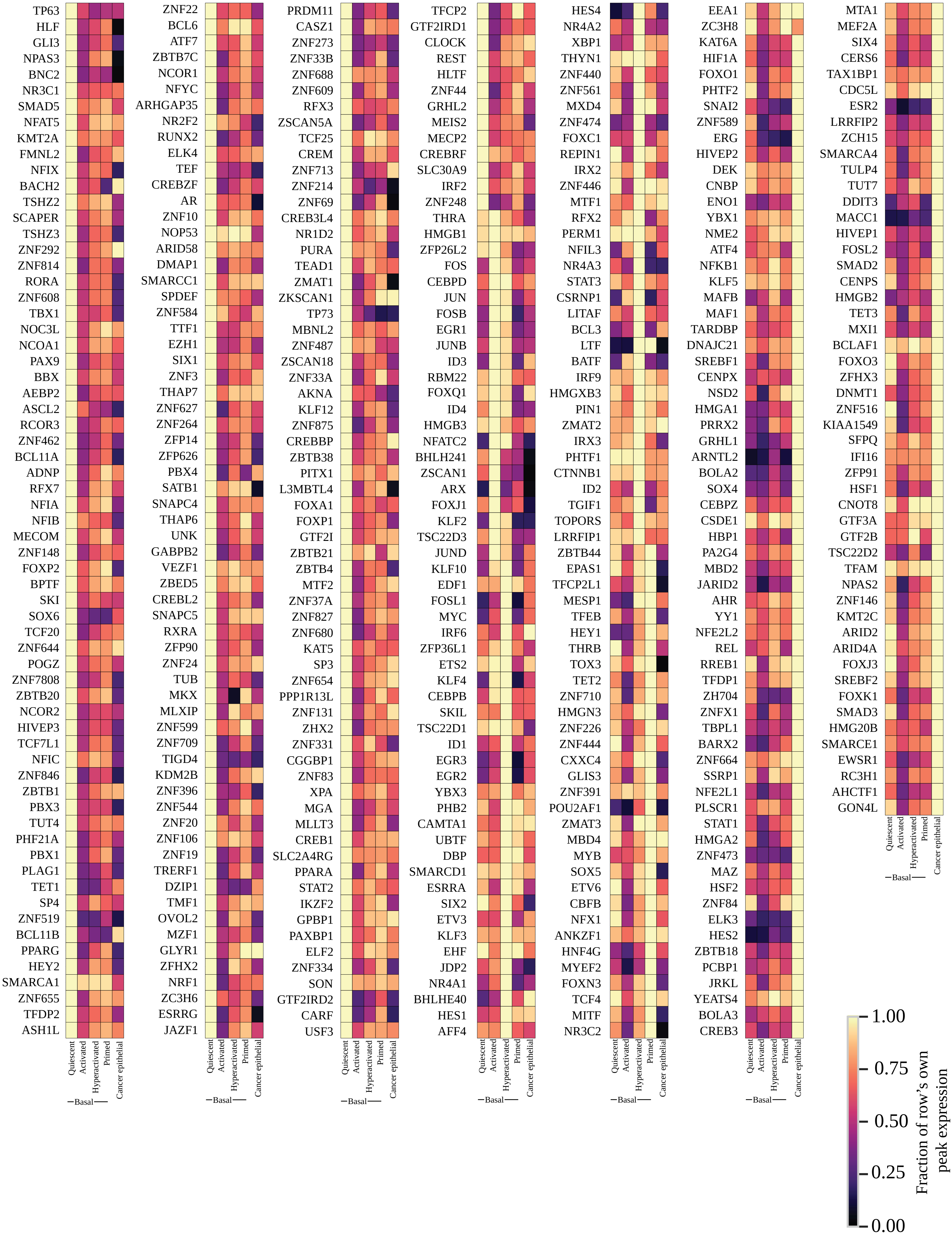


**Supplementary Figure E15**

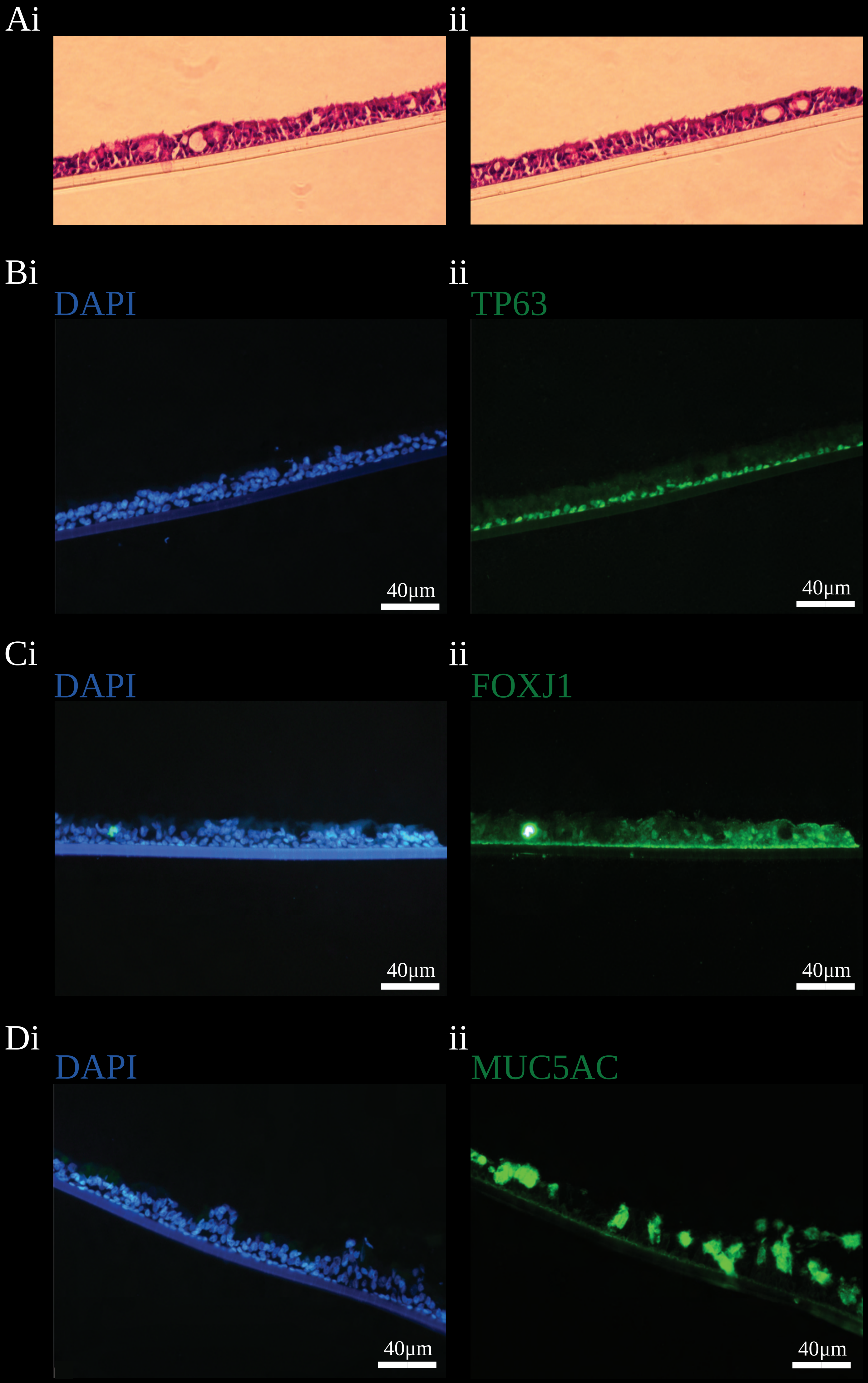

Ai
ii
Bi
DAPI
40µm
ii
TP63
40µm
Ci
DAPI
40µm
ii
FOXJ1
40µm
Di
DAPI
40µm
ii
MUC5AC
40µm

### unsorted eGFP negative cells

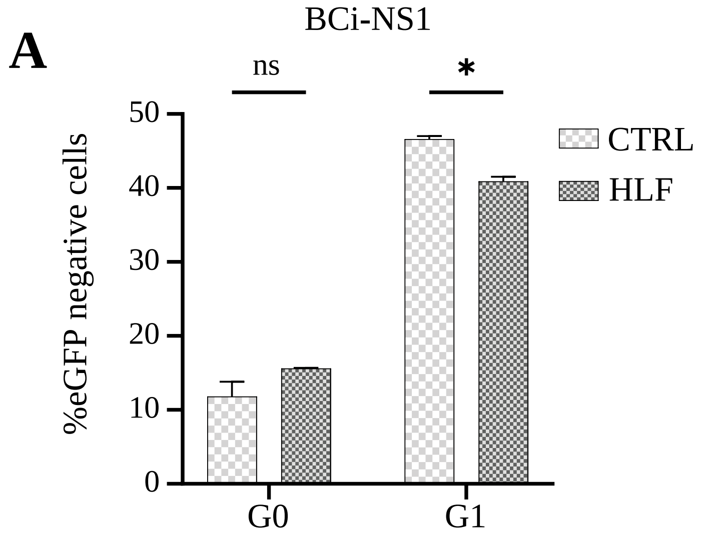


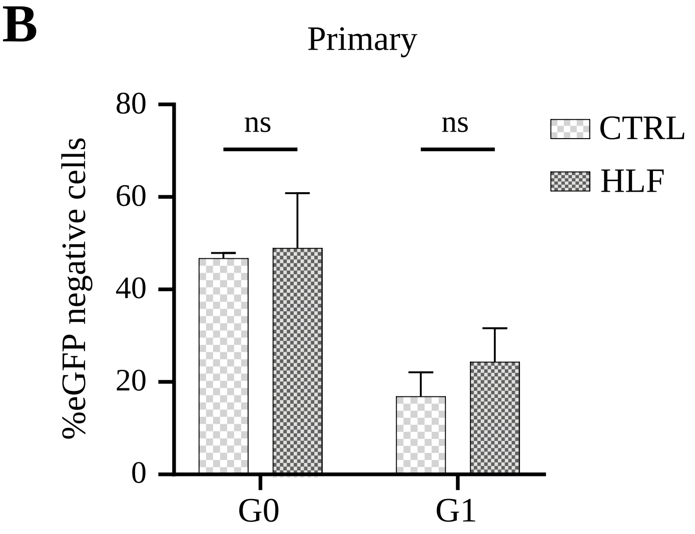


### unsorted eGFP positive cells

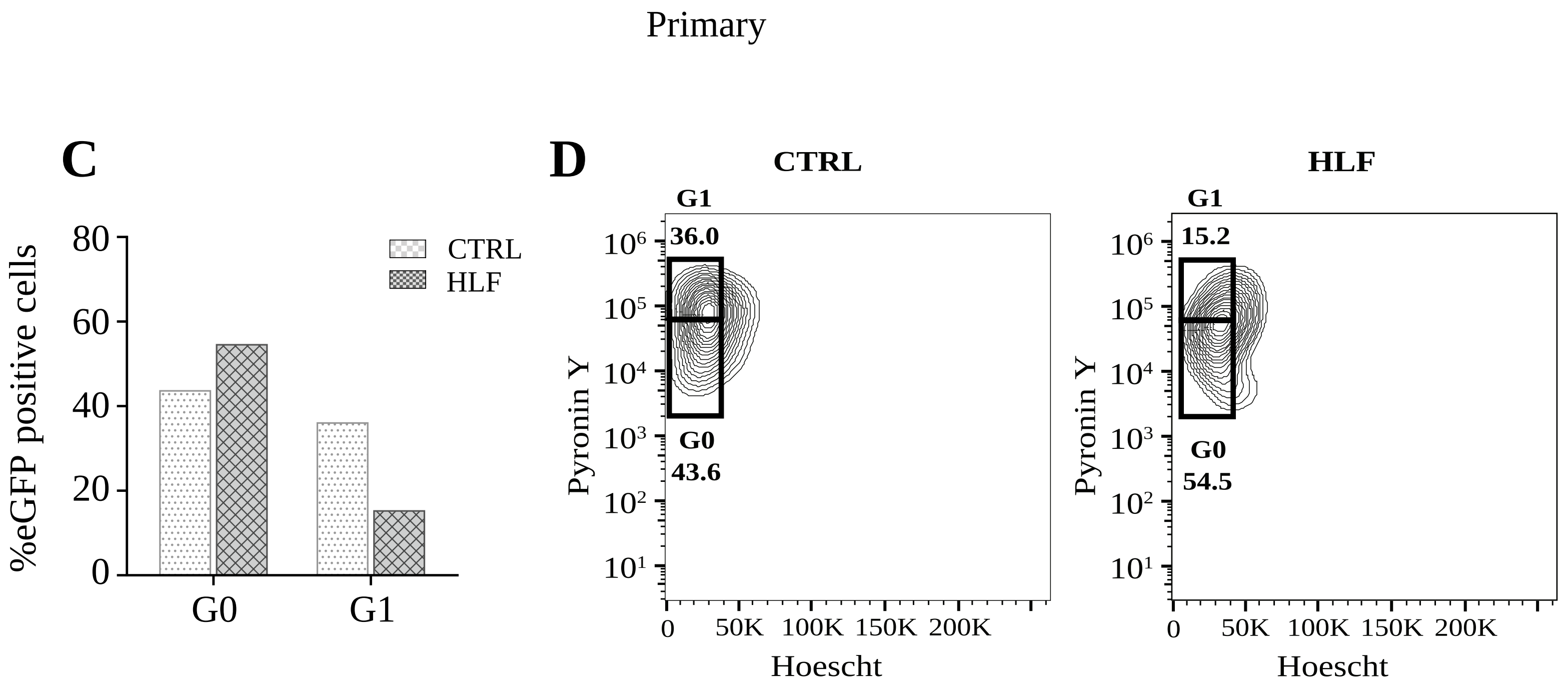


**Supplementary Figure E17**

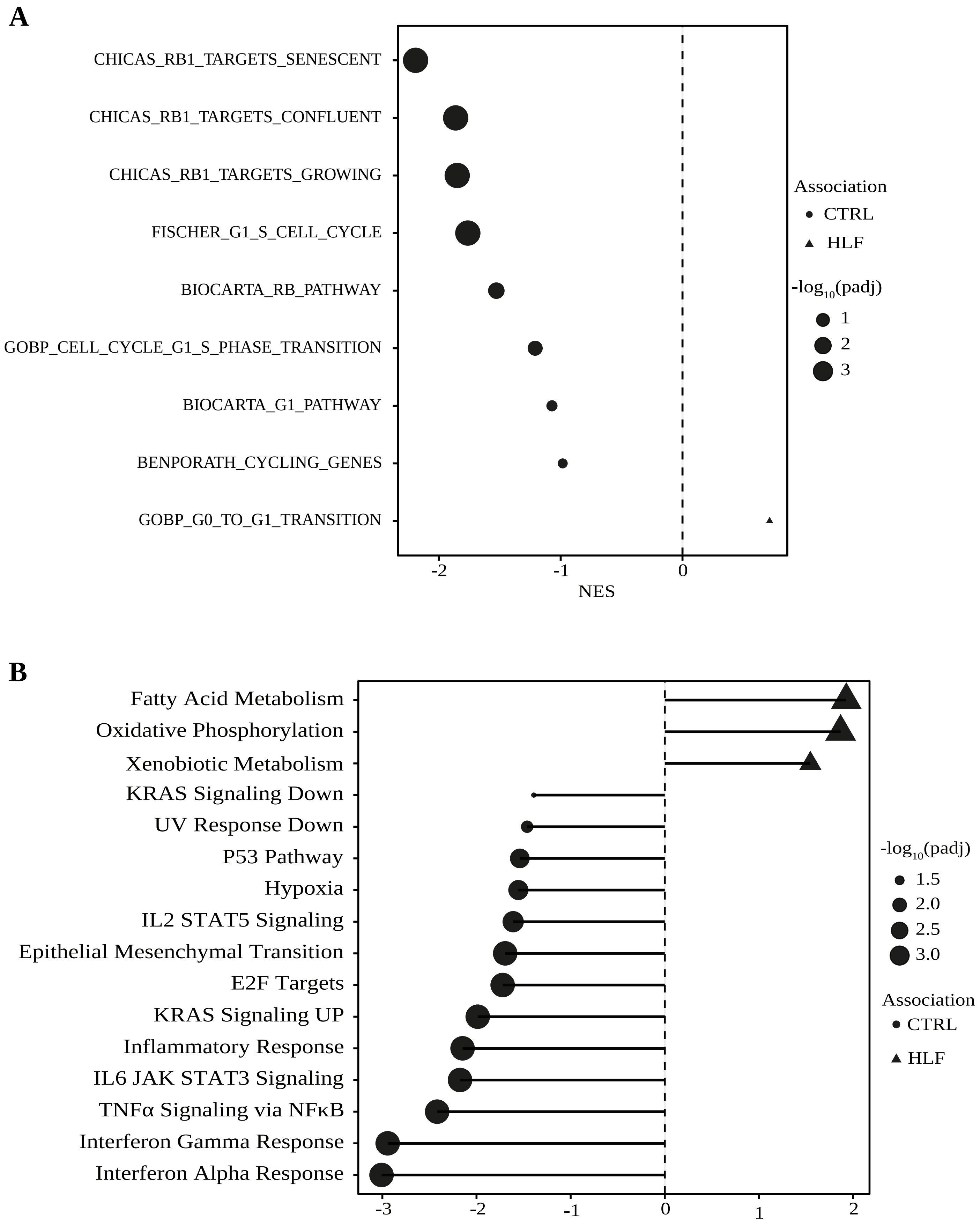


**Supplementary Figure E18**

### HLF pcr

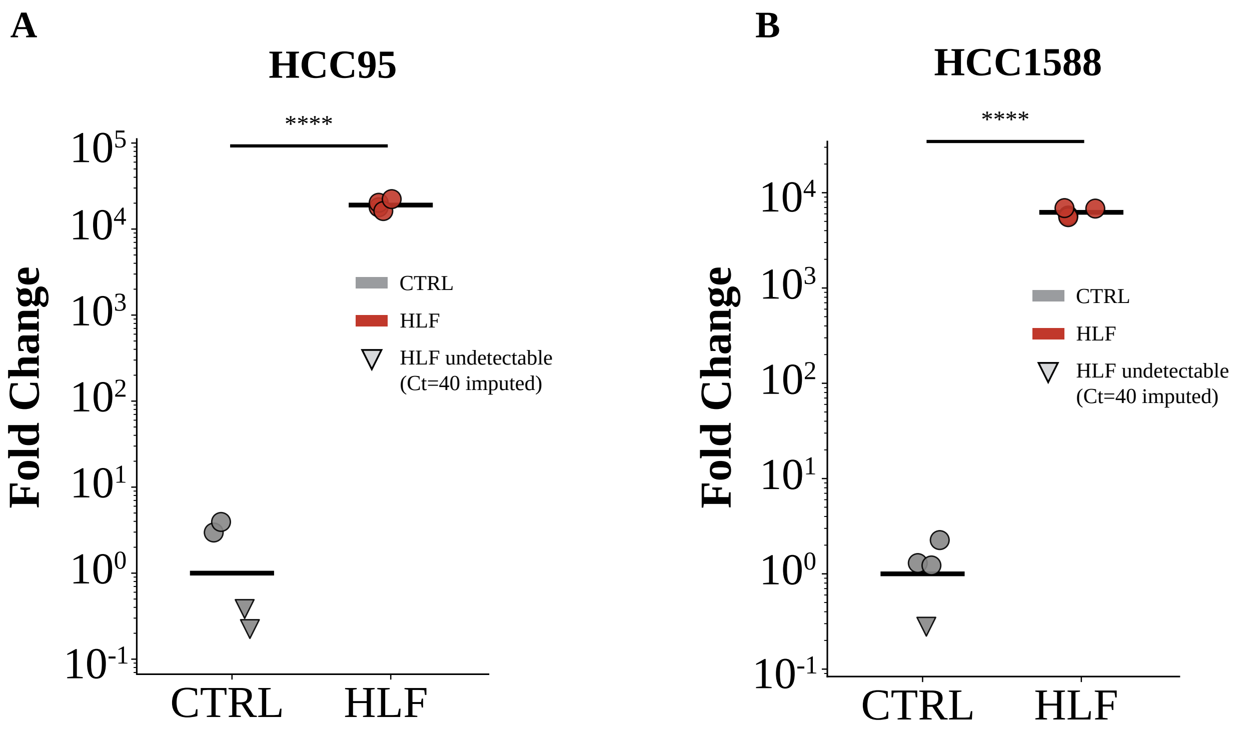


### HLF overexpression
### (ovx transduced, unsorted)

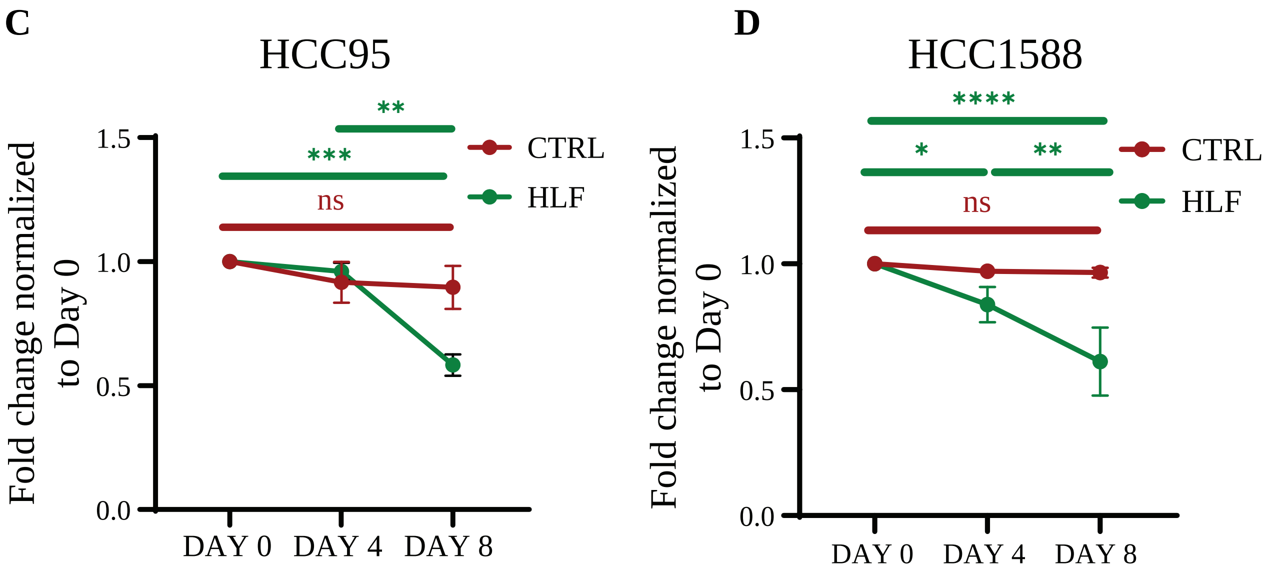


**Supplementary Figure E19**

**A**

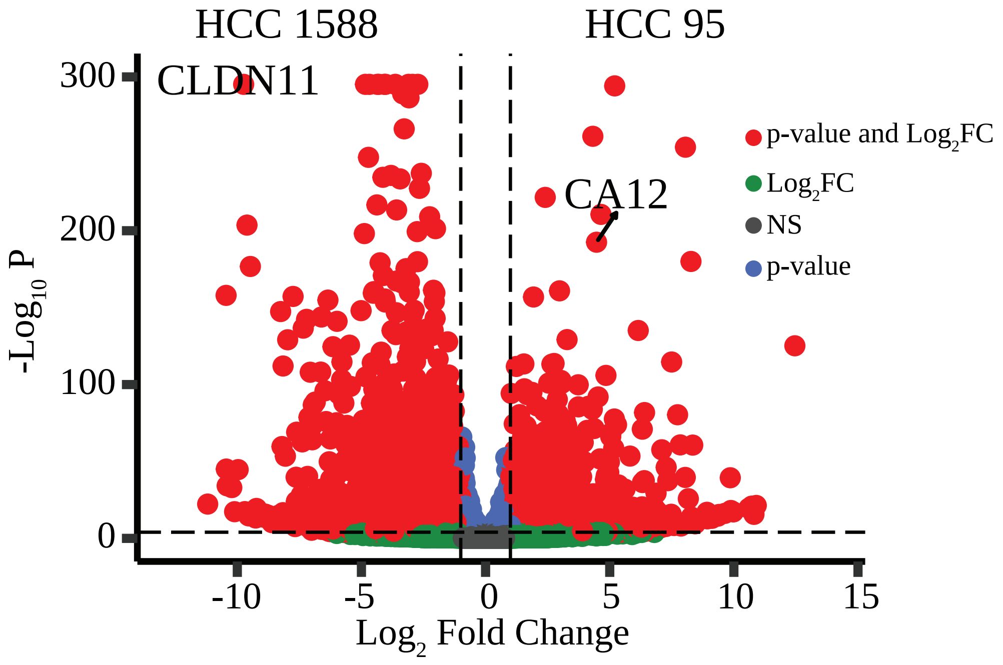


**B**

| | baseMean | log2FoldChange | lfcSE | stat | pvalue | padj |
|---|---|---|---|---|---|---|
| GCLC | 5906.377297 | -2.730517951 | 0.062611889 | -43.61021484 | 0 | 0 |
| CLDN11 | 3254.815505 | -9.766911552 | 0.22471737 | -43.46309122 | 0 | 0 |
| TIMP3 | 2317.220177 | -4.842957785 | 0.097379068 | -49.7330473 | 0 | 0 |
| CAT | 6902.987867 | -4.078482253 | 0.082315861 | -49.54673627 | 0 | 0 |
| TTC9 | 2644.27528 | -3.641162918 | 0.096647658 | -37.67461097 | 0 | 0 |
| EHF | 15275.43273 | -4.684597488 | 0.093966473 | -49.8539248 | 0 | 0 |
| GLUL | 38061.70393 | -2.943505446 | 0.056372266 | -52.21548906 | 0 | 0 |
| ITM2B | 15243.04316 | -3.102078445 | 0.079794562 | -38.87581261 | 0 | 0 |
| ABTB2 | 2256.076157 | -4.036041129 | 0.08005759 | -50.41422205 | 0 | 0 |
| TACSTD2 | 26456.70785 | -4.305088475 | 0.090728743 | -47.45010606 | 0 | 0 |
| HLA-B | 29978.31854 | -4.363254238 | 0.09151057 | -47.68033051 | 0 | 0 |
| FBN2 | 6070.141885 | 5.213273044 | 0.1420429 | 36.7021022 | 6.76E-295 | 1.45E-291 |
| CDC42EP3 | 4038.519943 | -3.342010935 | 0.091866108 | -36.37915018 | 9.10E-290 | 1.80E-286 |
| CASP4 | 2090.209325 | -3.08664058 | 0.085239364 | -36.21144553 | 4.02E-287 | 7.40E-284 |
| PDHX | 4438.707027 | -3.280586022 | 0.093981394 | -34.90676054 | 5.87E-267 | 1.01E-263 |
| UGT1A7 | 713.7840212 | 4.323157429 | 0.124993625 | 34.58702342 | 3.96E-262 | 6.37E-259 |
| MTND1P2 | 1074.556732 | 8.097092708 | 0.23737083 | 34.11157435 | 4.97E-255 | 7.53E-252 |
| SLPI | 4482.468654 | -4.726003371 | 0.140385682 | -33.66442581 | 1.92E-248 | 2.74E-245 |
| CYB561A3 | 3013.879223 | -2.589681284 | 0.078601363 | -32.94702752 | 4.67E-238 | 6.32E-235 |
| AMIGO2 | 763.1433635 | -3.815624004 | 0.116145418 | -32.85212692 | 1.06E-236 | 1.37E-233 |
| TRIM44 | 17533.69247 | -4.142163027 | 0.126415638 | -32.76622327 | 1.78E-235 | 2.19E-232 |
| KYNU | 2918.663576 | -3.447231208 | 0.105443702 | -32.692623 | 1.99E-234 | 2.33E-231 |
| HS6ST1 | 5529.096659 | -2.663833006 | 0.082579286 | -32.25788368 | 2.73E-228 | 3.05E-225 |
| KRT14 | 13085.02666 | 2.404670533 | 0.075534769 | 31.83528035 | 2.10E-222 | 2.26E-219 |
| APIP | 2483.808147 | -4.385896165 | 0.139321007 | -31.48050858 | 1.61E-217 | 1.65E-214 |
| SLC37A2 | 2024.484797 | -3.588250883 | 0.114863837 | -31.23916966 | 3.13E-214 | 3.10E-211 |
| MTCO1P1 | 9051.253859 | 4.659702538 | 0.15018507 | 31.02640321 | 2.37E-211 | 2.27E-208 |
| HLA-A | 28797.57351 | -2.254127373 | 0.07292142 | -30.91173186 | 8.31E-210 | 7.64E-207 |
| GFRA1 | 1392.555148 | -9.638278089 | 0.315952913 | -30.50542566 | 2.21E-204 | 1.96E-201 |
| CUX1 | 5540.26909 | -2.017977066 | 0.066532823 | -30.33054938 | 4.54E-202 | 3.89E-199 |
| ARPIN | 1056.924225 | -2.756523754 | 0.091320973 | -30.18500212 | 3.73E-200 | 3.10E-197 |
| TMEM156 | 963.6317887 | -4.894119157 | 0.162666129 | -30.08689747 | 7.19E-199 | 5.79E-196 |
| CA12 | 1462.704981 | 4.479334699 | 0.151049603 | 29.65472669 | 2.95E-193 | 2.30E-190 |
| ESRG | 665.1275259 | 8.276945769 | 0.288707379 | 28.66897898 | 9.30E-181 | 7.04E-178 |
| SCPEP1 | 6018.019925 | -2.744857264 | 0.095786314 | -28.65604842 | 1.35E-180 | 9.92E-178 |
| SCNN1A | 8018.126722 | -4.258804463 | 0.148914367 | -28.59901674 | 6.91E-180 | 4.94E-177 |
| CP | 2411.690705 | -9.614297196 | 0.338454594 | -28.40646095 | 1.68E-177 | 1.17E-174 |
| B2M | 29544.05832 | -3.206998068 | 0.113360581 | -28.29024014 | 4.56E-176 | 3.09E-173 |
| GPR87 | 1037.598455 | -4.125720558 | 0.14773257 | -27.92695319 | 1.26E-171 | 8.29E-169 |
| PDPN | 1017.524988 | -3.590755876 | 0.129972009 | -27.62714761 | 5.25E-168 | 3.38E-165 |
| CYP4F11 | 5784.46991 | -3.070537143 | 0.111368544 | -27.57095518 | 2.48E-167 | 1.56E-164 |
| HLA-F | 2283.979357 | -3.381972638 | 0.122839154 | -27.53171556 | 7.33E-167 | 4.49E-164 |
| CD24 | 18272.7143 | -2.09914556 | 0.077387324 | -27.12518611 | 4.97E-162 | 2.98E-159 |
| MARCHF4 | 525.442029 | 2.980903596 | 0.110026283 | 27.09265018 | 1.20E-161 | 7.04E-159 |
| LINC0150[illegible] | 437.4679251 | -4.505973964 | 0.166588829 | -27.04847619 | 3.98E-161 | 2.28E-158 |
| HLA-C | 20022.87732 | -3.077836453 | 0.113912607 | -27.01927842 | 8.77E-161 | 4.91E-158 |
| TSKU | 4254.171008 | -2.042119301 | 0.075692268 | -26.97923267 | 2.59E-160 | 1.42E-157 |
| UPK3B | 576.237249 | -4.533929501 | 0.168165371 | -26.96113644 | 4.22E-160 | 2.27E-157 |
| WFDC2 | 2976.917711 | -10.64094903 | 0.396427917 | -26.8420779 | 1.04E-158 | 5.49E-156 |
| AFAP1-AS[illegible] | 655.7279387 | -7.836752424 | 0.292607899 | -26.78243633 | 5.18E-158 | 2.67E-155 |

**Supplementary Figure E20**

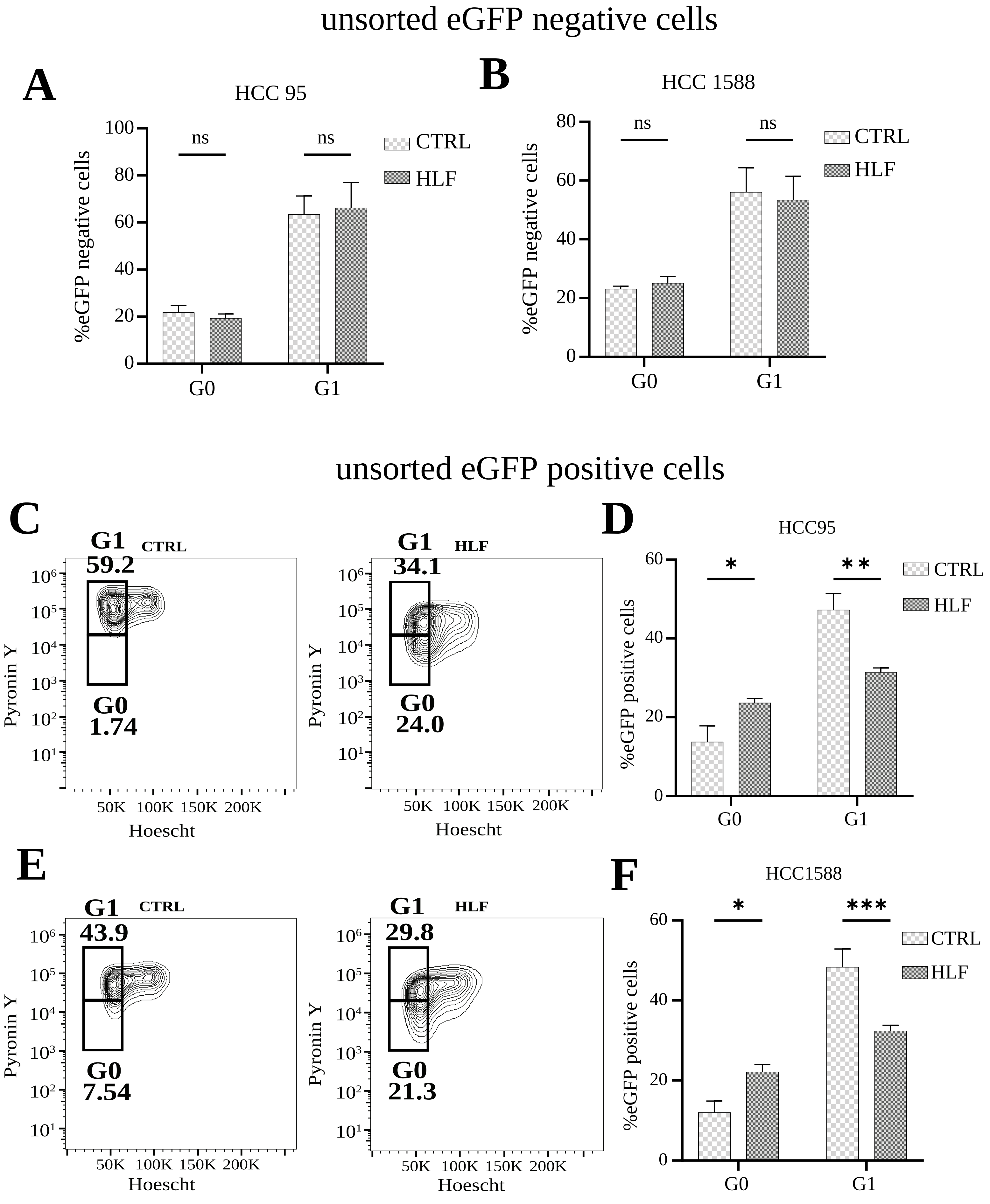


**Supplementary Figure E21**

Primary Cancer organoids

CTRL DAPI

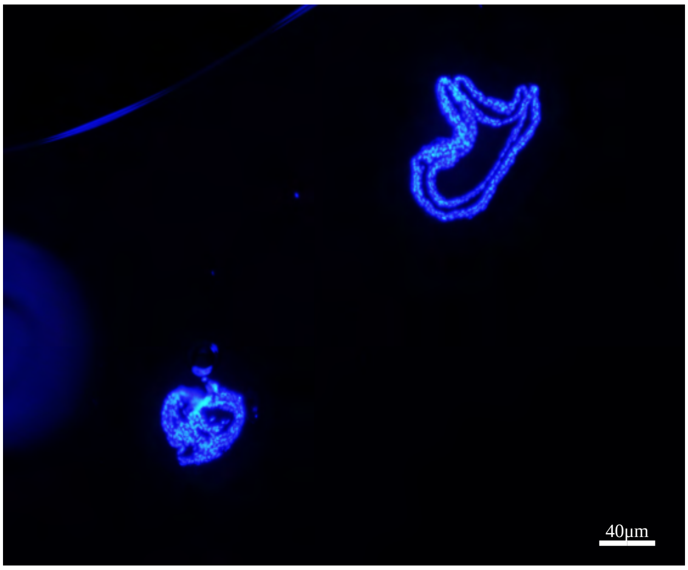


HLF DAPI

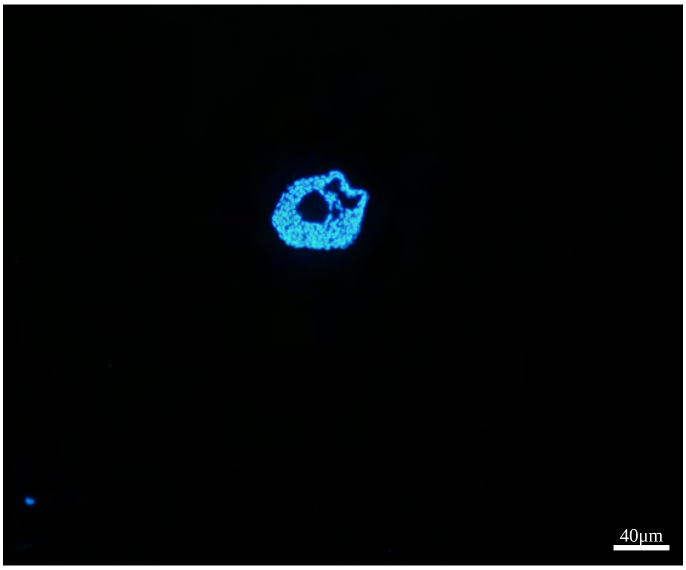


CTRL TP63

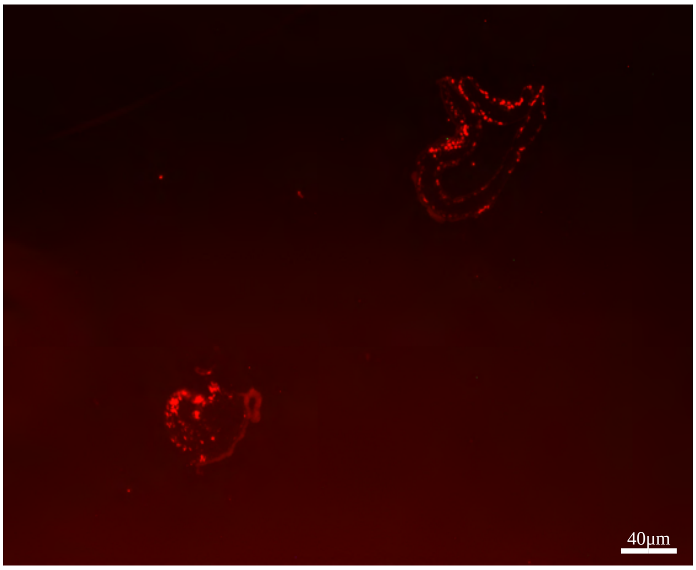


HLF TP63

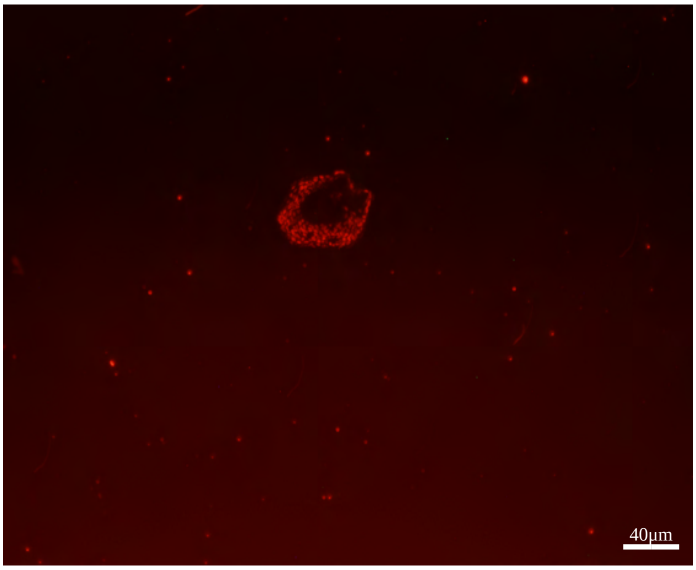


CTRL eGFP

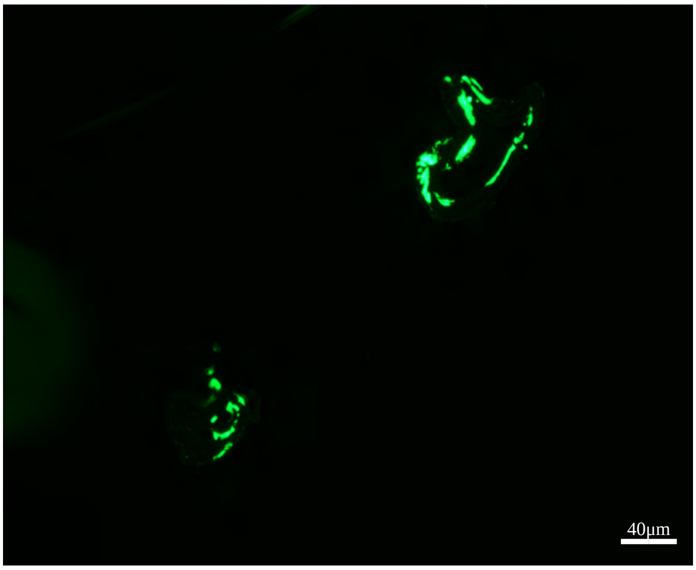


HLF overexpression eGFP

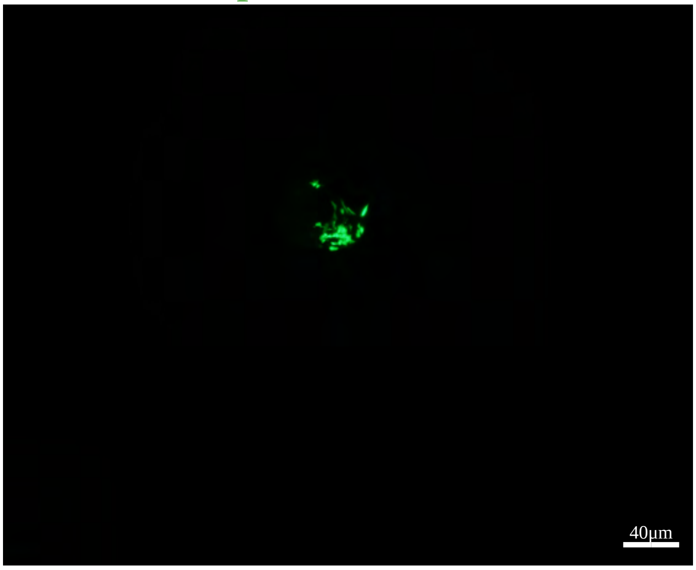


Supplementary Figure E22
