## Supplementary table 1 for "HLF suppresses DREAM target genes and reinstates quiescence in lung squamous cell carcinoma"

| | Sample ID | Year of Birth | Age | Age Classification | Sex | Smoking Status | Stage of tumor | Fusions or Amplifications | Mutation | Staining |
|---|---|---|---|---|---|---|---|---|---|---|
| Healthy | | | | | | | | | | |
| | Healthy 1 | N/A | N/A | N/A | N/A | Healthy never smoker | | | | |
| | Healthy 2 | 1996 | 23 | Young | M | Healthy never smoker | | | | |
| | Healthy 3 | 1989 | 29 | Young | F | Healthy never smoker | | | | |
| | Healthy 4 | 1979 | 39 | Young | M | Healthy never smoker | | | | |
| | Healthy 5 | 1998 | 20 | Young | F | Healthy never smoker | | | | |
| | Healthy 6 | 1953 | 66 | Aged | M | Healthy never smoker | | | | |
| | Healthy 7 | 1955 | 64 | Aged | F | Healthy never smoker | | | | |
| | Healthy 8 | 1984 | 35 | Young | F | Healthy never smoker | | | | |
| | Healthy 9 | 1944 | 75 | Aged | F | Healthy never smoker | | | | |
| | Healthy 10 | 1996 | 22 | Young | F | Healthy never smoker | | | | |
| Cancer | Cancer 1 | 1945 | 81 | Aged | M | Smoker | pT2b N0 (IIA) | NOT DETECTED: EGFR, BRAF, ALK, ROS1, KRAS, RET fusion | Unknown | $CK5^{+ive}$, $p40^{+ive}$, $TTF^{-ive}$ |
| | Cancer 2 | 1961 | 65 | Aged | M | Smoker | pT4 N0 (IIIA) | NOT DETECTED: EGFR, BRAF, ALK, ROS1, KRAS, RET fusion | Unknown | $CK5^{+ive}$, $p40^{+ive}$, $TTF^{-ive}$ |
| | Cancer 3 | 1951 | 75 | Aged | F | Smoker | pT2a N0 (IB) | NOT DETECTED: EGFR, BRAF, ALK, ROS1, KRAS, RET fusion. FGFR mutation detected | FGFR3 | $CK5^{+ive}$, $p40^{+ive}$, $TTF^{-ive}$ |
| | Cancer 4 | 1958 | 68 | Aged | M | Smoker | pT3N1 | NOT DETECTED: EGFR, BRAF, ALK, ROS1, KRAS, RET fusion | Unknown | $CK5^{+ive}$, $p40^{+ive}$, $TTF^{-ive}$ |
| | Cancer 5 | 1961 | 65 | Aged | F | Smoker | pT2aN0 | NOT DETECTED: EGFR, BRAF, ALK, ROS1, KRAS, RET fusion | Unknown | $CK5^{+ive}$, $p40^{+ive}$, $TTF^{-ive}$ |
| | Cancer 6 | 1950 | 76 | Aged | M | Smoker | pT3N2 PL3 | NOT DETECTED: EGFR, BRAF, ALK, ROS1, KRAS, RET fusion | Unknown | $CK5^{+ive}$, $p40^{+ive}$, $TTF^{-ive}$ |

Supplementary Table 1
