## Supplementary table 8 for "HLF suppresses DREAM target genes and reinstates quiescence in lung squamous cell carcinoma"

| Basal I | Basal I (extended) | Basal II | Basal III | Basal IV |
|---|---|---|---|---|
| GLI3 | ZNF655 | HMGB1 | KLF10 | UBTF |
| NPAS3 | TFDP2 | DEK | ENO1 | MAFB |
| KAT6A | ASH1L | EPAS1 | FOS | ID3 |
| BNC2 | ZNF22 | NOP53 | EDF1 | RBM22 |
| NR3C1 | BCL6 | CNBP | CEBPD | FOXQ1 |
| SMAD5 | ATF7 | ARID5B | FOSL1 | DBP |
| NFAT5 | ZBTB7C | ZFP36L2 | MYC | MAF1 |
| KMT2A | NCOR1 | | JUN | SMARCD1 |
| FMNL2 | HIVEP2 | | YBX1 | ID4 |
| HIF1A | NFYC | | IRF6 | ESRRA |
| TP63 | ARHGAP35 | | ZFP36L1 | TARDBP |
| NFIX | NR2F2 | | NME2 | DNAJC21 |
| BACH2 | RUNX2 | | FOSB | SREBF1 |
| TSHZ2 | ELK4 | | EGR1 | TFCP2L1 |
| SCAPER | ZBTB44 | | ETS2 | SIX2 |
| TSHZ3 | TEF | | KLF4 | CENPX |
| ZNF292 | THRA | | ATF4 | NSD2 |
| ZNF814 | CREBZF | | CEBPB | HMGA1 |
| RORA | AR | | SKIL | MESP1 |
| FOXO1 | ZNF10 | | JUNB | ETV3 |
| ZNF608 | | | TSC22D1 | TFEB |
| TBX1 | | | ID1 | DMAP1 |
| NOC3L | | | EGr3 | PRRX2 |
| NCOA1 | | | EGR2 | GRHL1 |
| PAX9 | | | NFKB1 | ARNTL2 |
| BBX | | | YBX3 | BOLA2 |
| AEBP2 | | | PHB2 | KLF3 |
| ASCL2 | | | CAMTA1 | SOX4 |
| HLF | | | KLF5 | EHF |
| ZNF538 | | | | CEBPZ |
| RCOR3 | | | | HMGB3 |
| ZNF462 | | | | CSDE1 |
| BCL11A | | | | JDP2 |
| ADNP | | | | NR4A1 |
| RFX7 | | | | HBP1 |
| NFIA | | | | PA2G4 |
| NFIB | | | | MBD2 |
| MECOM | | | | JARID2 |
| ZNF148 | | | | AHR |
| FOXP2 | | | | BHLHE40 |
| BPTF | | | | HES1 |
| SKI | | | | HEY1 |
| PHTF2 | | | | SMARCC1 |
| SOX6 | | | | SKIL |
| TCF20 | | | | AFF4 |
| ZNF644 | | | | HES4 |
| POGZ | | | | YY1 |
| ZNF780B | | | | NFE2L2 |
| ZBTB20 | | | | REL |
| NCOR2 | | | | |
| HIVEP3 | | | | |
| TCF7L1 | | | | |
| NFIC | | | | |
| ZNF846 | | | | |
| ZBTB1 | | | | |
| PBX3 | | | | |
| TUT4 | | | | |
| PHF21A | | | | |
| PBX1 | | | | |
| PLAG1 | | | | |
| TET1 | | | | |
| SNAI2 | | | | |
| ZNF589 | | | | |
| SP4 | | | | |
| ZNF519 | | | | |
| BCL11B | | | | |
| ERG | | | | |
| PPARG | | | | |
| HEY2 | | | | |
| SMARCA1 | | | | |

**Supplementary Table 7**
