## Supplementary table 2 for "HLF suppresses DREAM target genes and reinstates quiescence in lung squamous cell carcinoma"

| SampleID | Year of birth | Age | Age classification | Year biopsy taken | Gender | Smoking status |
|---|---|---|---|---|---|---|
| Healthy 1 | N/A | N/A | N/A | N/A | N/A | Healthy never smoker |
| Healthy 2 | 1996 | 23 | Young | 2019 | M | Healthy never smoker |
| Healthy 3 | 1989 | 29 | Young | 2019 | F | Healthy never smoker |
| Healthy 4 | 1979 | 39 | Young | 2019 | M | Healthy never smoker |
| Healthy 5 | 1998 | 20 | Young | 2019 | F | Healthy never smoker |
| Healthy 6 | 1953 | 66 | Aged | 2019 | M | Healthy never smoker |
| Healthy 7 | 1955 | 64 | Aged | 2019 | F | Healthy never smoker |
| Healthy 8 | 1984 | 35 | Young | 2019 | F | Healthy never smoker |
| Healthy 9 | 1944 | 75 | Aged | 2019 | F | Healthy never smoker |
| Healthy 10 | 1996 | 22 | Young | 2019 | F | Healthy never smoker |

| **Sample name** | **FEV1** | **FEV1 % predicted** | **FEV1/ FVC** | **DLCO % predicted** | **Leukocytes** | **Neutrophils** | **Eosinophils** | **Basophils** | **Lymphocytes** | **Monocytes** |
|---|---|---|---|---|---|---|---|---|---|---|
| Healthy 1 | N/A | | | | | | | | | |
| Healthy 2 | 3.49 | 101 | 0.87 | 104 | 5.6 | 2.1 | 0.3 | <0.1 | 2.7 | 0.3 |
| Healthy 3 | 4.07 | 111 | 0.81 | 92 | 7.8 | 3.2 | 0.2 | <0.1 | 4.2 | 0.2 |
| Healthy 4 | 4.75 | 104 | 0.83 | 106 | 4.3 | 2.2 | 0.1 | <0.1 | 1.7 | 0.3 |
| Healthy 5 | 3.22 | 87 | 0.92 | 74 | 6.1 | 3.4 | 0.3 | <0.1 | 1.9 | 0.5 |
| Healthy 6 | 3.51 | 122 | 0.81 | 81 | 3.5 | 1.7 | 0.1 | <0.1 | 1.5 | 0.2 |
| Healthy 7 | 4.23 | 107 | 0.77 | 96 | 7.7 | 4.5 | 0.1 | <0.1 | 2.5 | 0.5 |
| Healthy 8 | 3.18 | 98 | 0.81 | 95 | 6.4 | 4.0 | 0.1 | <0.1 | 1.9 | 0.5 |
| Healthy 9 | 1.91 | 85 | 0.67 | 62 | 6.9 | 2.8 | 0.2 | <0.1 | 3.4 | 0.5 |
| Healthy 10 | 4.12 | 118 | 0.98 | 107 | 6.5 | 4.4 | 0.1 | <0.1 | 1.6 | 0.4 |

Graphical representation of the Clinical characteristics of the patient samples

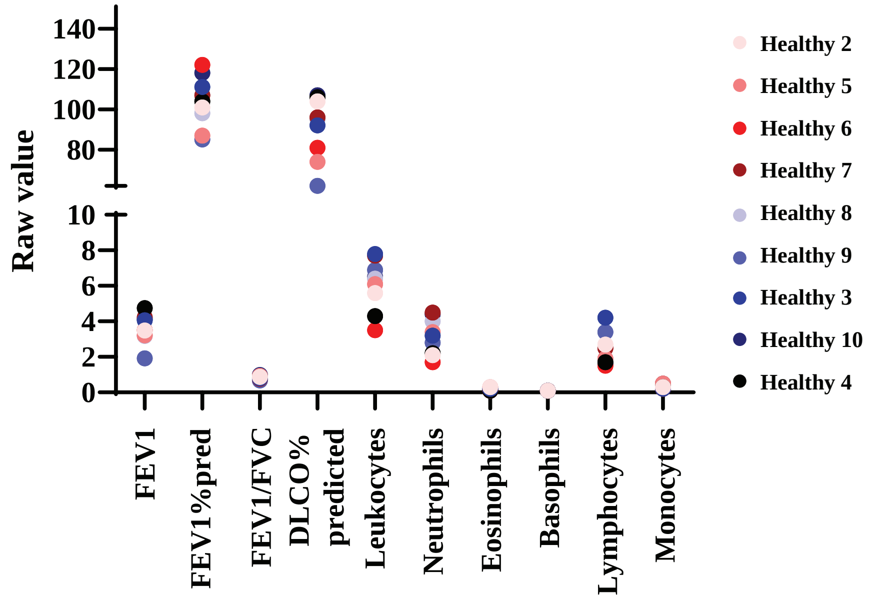

Supplementary Table 2
